# PI3K signaling promotes inflammatory tumor-macrophage crosstalk associated with mesenchymal glioblastoma

**DOI:** 10.64898/2026.07.27.738732

**Authors:** Rebeca Burgos-Panadero, María Rosado-Sanz, Victor Montosa-i-Micó, Zoey J Tolboom, Nuria Martínez-Alarcón, Macarena Ferrero, Eloi Casals, Anna Esteve-Codina, Juan M Garcia-Gomez, Michaël H Meel, Jaime Font de Mora

## Abstract

**Background:** Glioblastoma (GBM) is a highly heterogeneous and vascularized malignancy in which the mesenchymal (MES) subtype is associated with poor prognosis, extensive macrophage infiltration and resistance to therapy. However, the signaling mechanisms integrating vascular remodeling with inflammatory tumor-macrophage crosstalk remain incompletely understood.

**Methods:** We integrated magnetic resonance imaging-derived vascular phenotyping with transcriptomic analyses of human glioblastoma cohorts to identify molecular pathways associated with highly vascular tumors. Functional studies using glioblastoma cell lines, THP-1-derived macrophages and co-culture systems were performed to investigate the role of PI3K signaling in tumor-macrophage communication. Finally, an independent single-cell transcriptomic cohort of primary human glioblastoma was interrogated to determine whether the identified inflammatory programs were conserved in malignant cells from patient tumors.

**Results:** Integrated imaging-transcriptomic analyses identified highly vascular glioblastomas as tumors enriched for the MES subtype, increased macrophage infiltration and activation of PI3K-associated signaling. Pharmacological inhibition of PI3K reduced the expression of macrophage-recruiting cytokines and impaired the ability of glioblastoma cells to educate macrophages toward an immunosuppressive phenotype. Reciprocally, tumor-educated macrophages enhanced inflammatory signaling, immune checkpoint expression and migratory capacity in glioblastoma cells, whereas IL-6 blockade attenuated these effects, identifying IL-6 as a key mediator of this bidirectional communication. To determine whether these inflammatory programs were conserved in human disease, we analyzed an independent single-cell transcriptomic dataset of primary glioblastomas. MES-like malignant cells exhibited the strongest inflammatory transcriptional programs among the four malignant transcriptional states, including higher NF-κB activation program scores and tumor-macrophage communication signature scores. At the tumor level, MES-like enrichment was positively associated with higher inflammatory program activity, supporting the clinical relevance of the proposed signaling axis.

**Conclusions:** Together, our findings identify PI3K signaling as a central regulator integrating vascular remodeling with inflammatory tumor-macrophage communication in mesenchymal glioblastoma. These results provide a mechanistic framework linking PI3K signaling, macrophage education and the MES phenotype, and provide a rationale for therapeutic strategies aimed at disrupting inflammatory signaling within the glioblastoma microenvironment.

## INTRODUCTION

Glioblastoma (GBM) is the most prevalent primary brain tumor in adults, characterized by its high aggressiveness, with a median survival of only 15 months and a 5-year survival rate of approximately 7% despite aggressive treatment regimens [1, 2]. While diagnostic and prognostic markers, such as microvascular proliferation, necrosis, and molecular features including *IDH* mutations, 1p/19q codeletion, *MGMT* promoter methylation, and *EGFRvIII* amplification, are routinely assessed and have refined disease classification, therapeutic strategies directed at these molecular alterations (including IDH1 inhibitors and EGFRvIII-targeted approaches) have thus far failed to produce meaningful clinical benefit, and overall patient outcomes remain poor [3]. Consequently, radiomic, genomic, transcriptomic, and epigenetic insights are crucial for enhancing the diagnosis, prognosis, and therapeutic strategies for GBM patients.

Molecular classifications have provided essential frameworks for identifying therapeutic targets. The distinction between primary GBM (IDH wild-type) and secondary GBM (*IDH* mutant) reflects differences in prognosis, with the former accounting for 76% of GBM cases and exhibiting less favorable outcomes [4, 5]. Common genomic alterations in IDH wild-type GBM include *PTEN* mutations, *EGFR* amplification, and chromosome 10 loss. Conversely, IDH-mutant GBM frequently harbors *ATRX* with fewer *PTEN* and *EGFR* alterations [6–8]. Additionally, gene-expression subtypes (proneural, mesenchymal, and classical) delineate distinct molecular pathways associated with tumor progression. Specifically, the mesenchymal subtype, which displays the poorest prognosis and strongest therapy resistance and is frequently associated with *PTEN* loss, shows activation of the PI3K/AKT pathway, contributing to immune suppression and tumor proliferation [9].

Despite these classifications, GBM remains a highly heterogeneous tumor with near-universal recurrence after treatment. This underscores the urgent need for multimodal therapeutic strategies that integrate novel approaches such as chemotherapy, anti-angiogenic agents, radiotherapy, and immunotherapy [10, 11]. A critical barrier to progress is the complex interaction between tumor cells and the surrounding tumor microenvironment (TME), which comprises immune cells, blood vessels, and stromal elements. The TME in GBM is predominantly immunosuppressive, with high infiltration of microglia, and tumor-associated macrophages (TAMs), collectively accounting for up to 50% of the tumor mass [12, 13]. Notably, TAMs are strongly associated with the mesenchymal subtype [14].

Genomic studies have revealed that specific alterations, such as *PTEN* loss, promote an immune landscape dominated by M2-like macrophages, myeloid-derived suppressor cells (MDSCs), and regulatory T cells (Tregs), exacerbating immunosuppression [15]. Concurrently, the vascular niche facilitates the delivery of oxygen and nutrients to the tumor, compounding its aggressive phenotype.

Advanced imaging techniques, such as ONCOhabitats-based segmentation of magnetic resonance imaging (MRI), have revealed vascular heterogeneity within the tumor [16, 17]. This variability correlates with patient survival and response to therapies like bevacizumab and temozolomide [18, 19], and it has also demonstrated a dependence/correlation between specific vascular patterns and *IDH* status [20].

In this study, we investigate how tumor-intrinsic PI3K signaling shapes the immunosuppressive macrophage microenvironment in GBM. Using ONCOhabitats software to stratify tumors by MRI-derived vascular biomarkers such as relative cerebral blood volume (rCBV), we show that highly vascularized GBMs are enriched for PI3K pathway alterations and increased macrophage infiltration. Mechanistically, we demonstrate that PI3K-active GBM cells promote an M2-like macrophage phenotype through cytokine-dependent paracrine signaling involving IL-6 and IL-10, and that IL-6 blockade disrupts this immunoregulatory crosstalk. These findings suggest that PI3K-dependent vascular phenotypes in GBM drive macrophage polarization and identify PI3K and IL-6 inhibitors as rational complementary strategies to overcome immunosuppression.

## MATERIALS AND METHODS

### Patient cohort

GBM patient data were obtained from The Cancer Genome Atlas glioblastoma project (TCGA-GBM) ([21]; available at the Genomic Data Commons: https://portal.gdc.cancer.gov/projects/TCGA-GBM), which comprises molecular and clinical data from 617 glioblastoma cases. For molecular subtype and immune deconvolution analyses, we included only tumors with available RNA-sequencing data. After exclusion of cases lacking RNA-seq data and those harboring IDH mutations, a total of 130 IDH-wild-type glioblastoma samples (we excluded 17 that lacked RNA-seq), were retained for transcriptomic analyses. For integrated radiogenomic analyses using ONCOhabitats ([16]; www.oncohabitats.upv.es), additional inclusion criteria were applied, requiring the availability of matched RNA-seq data, pre-operative MRI studies, and primary (non-recurrent) tumor samples. Fifteen patients met all criteria and were included in the imaging-transcriptomics cohort. MRI data were retrieved from The Cancer Imaging Archive (TCIA): https://wiki.cancerimagingarchive.net/pages/viewpage.action?pageId=1966258.

### TCGA GBM molecular subtype classification and pathway annotation

TCGA GBM samples (IDH-wild type) were classified into proneural, mesenchymal, and classical subtypes according to the Verhaak gene-expression signature [7] using Bioconductor scripts (https://bioconductor.org/packages/devel/bioc/vignettes/TCGAbiolinks/inst/doc/subtypes.html; gbm = curatedTCGAData(“GBM”, “RNASeq2GeneNorm”, dry.run=FALSE, version=“2.1.1”); table(gbm$Original.Subtype) (v.3.22)) in R (v4.5.0). Gene expression data were retrieved from the TCGA database (RNA-seq V2 RSEM normalized counts). PI3K-altered cases were defined by the presence of *PIK3CA* or *PIK3R1* mutations or *PTEN* mutations or deletions, as annotated in the TCGA mutation (MAF) and copy-number datasets. Immune cell composition was inferred using CIBERSORTx (https://cibersortx.stanford.edu/) with the LM22 signature matrix, 1 000 permutations, and quantile normalization disabled. Gene-set enrichment analyses were performed using the Parametric Analysis of Gene-set Enrichment (PAGE) algorithm [22] applied to KEGG pathways and the BROAD 2020.09 C2 curated gene collection and implemented in the R2 Genomics Analysis and Visualization Platform (http://r2.amc.nl). For the independent single-cell validation analyses, malignant-cell transcriptional programs were quantified using the published Neftel MES-like, AC-like, OPC-like and NPC-like gene signatures [23]. NF-κB activation was quantified using a curated NF-κB activation program gene signature (NFKBIA, TNFAIP3, RELB, BIRC3, TRAF1, TNIP1, BCL2A1, PLAU, ICAM1 and VCAM1). Tumor-macrophage communication was quantified using a curated tumor-macrophage communication signature (CCL2, CSF1, IL6, TGFB1, VEGFA, CD274, GAS6 and HGF).

### Single-cell transcriptomic analysis

An independent Smart-seq2 single-cell RNA-sequencing dataset of primary human glioblastomas generated by Neftel *et al.* [23] was used for validation analyses. Processed single-cell expression data were obtained from GEO (GSE131928; GSM3828672; log2(TPM/10 + 1)). Samples corresponding to different regions of the same tumor (MGH105A-D) were merged prior to downstream analyses and treated as a single biological specimen, yielding 7,930 cells across 28 tumors before malignant-cell filtering. A Seurat (v5.5.1) object was constructed from the log-normalized matrix without further normalization; the 2,000 most variable genes were identified, scaled, and used for principal-component analysis (30 components), and cells were embedded in two dimensions by UMAP (30 PCs). Non-malignant cells were identified according to the strategy described by Neftel et al. [23]. Cells expressing macrophage (CD14, AIF1, FCER1G, FCGR3A, TYROBP, CSF1R), T-cell (CD2, CD3D, CD3E, CD3G), or oligodendrocyte (MBP, TF, PLP1, MAG, MOG, CLDN11) marker genes were excluded. Following filtering, 6,860 malignant cells from 28 primary glioblastomas were retained and re-embedded (variable-feature selection, scaling, PCA and UMAP as above) for all downstream analyses (Supplementary Table 1).

Each malignant cell was scored for the published Neftel malignant transcriptional states (MES-like, AC-like, OPC-like, and NPC-like) using Seurat’s AddModuleScore function with 100 control genes and a fixed random seed. Cells were assigned to one of these four states based on the argmax of their control-corrected module scores. In addition, an NF-κB activation program score (NFKBIA, TNFAIP3, RELB, BIRC3, TRAF1, TNIP1, BCL2A1, PLAU, ICAM1, and VCAM1) and a tumor-macrophage communication signature score (CCL2, CSF1, IL6, TGFB1, VEGFA, CD274, GAS6, and HGF) were also calculated using Seurat’s AddModuleScore function with 100 control genes and a fixed random seed.

Within each tumor, the Spearman correlation coefficient (ρ) between the MES-like transcriptional program score and the tumor-macrophage communication signature score or the NF-κB transcriptional program score was computed across malignant cells. To account for the nesting of cells within patients, a linear mixed model was fitted (lme4/lmerTest) with the standardized signature score as outcome, the standardized MES-like transcriptional program score as fixed effect, and a by-patient random slope and intercept, and fixed-effect significance was assessed using Satterthwaite-approximated degrees of freedom. At the tumor level, the percentage of MES-like malignant cells per tumor was correlated with the mean tumor-macrophage communication signature and NF-κB scores (Spearman ρ, n = 28 tumors), and the same analysis was repeated for all four Neftel states.

### MRI data analysis and ONCOhabitats segmentation

Preoperative MRI data were collected from IDH-wild-type GBM patients, including pre-and post-gadolinium contrast agent-enhanced T1-weighted, fluid-attenuated inversion recovery (FLAIR) T2-weighted and dynamic susceptibility contrast (DSC) T2*-weighted perfusion sequences. Data scans were analyzed using ONCOhabitats (v1.1) to delineate tumor subregions (habitats) and quantify their vascular parameters. Specifically, relative cerebral blood volume (rCBV) values were derived from Hemodynamic Tissue Signature DSC perfusion maps from the High Angiogenic Tumor habitats (HAT) and normalized to contralateral white matter. Patients were classified as High Vascularity (HV) when rCBV HAT ≥ 5.9 and Moderate Vascularity (MV) when < 5.9 [24]. Kaplan-Meier survival analysis was performed using GraphPad Prism (v10.4.2).

### Cell lines and culture conditions

Human GBM cell lines U87MG (CVCL_GP63), LN-18 (CVCL_0392), and LN-229 (RRID:CVCL_0393), were maintained in DMEM supplemented with 10 % fetal bovine serum (FBS), 2 mM L-glutamine, and 1 % penicillin-streptomycin at 37 °C and 5 % CO₂. THP-1 monocytes were cultured in RPMI-1640 containing 10 % FBS and differentiated into macrophage-like M0 cells by exposure to 100 ng/ml phorbol 12-myristate 13-acetate (PMA) for 24 h. Once THP-1 cells were differentiated into M0 macrophages, they were cocultured with tumor cells for 72 h using 1 μm pore size inserts (Falcon 353095)[19]. Also, conditioned media (CM) from different and/or consecutive coculture experiments (as indicated) were isolated and used in 1:3 ratio with fresh media. CM were then centrifuged at 465xg for 5 minutes, and the supernatants were stored at-80 °C. Cells were regularly tested for mycoplasma contamination using the Mycoplasma PCR Detection Kit (Applied Biological Materials Inc.; G238). Cell lines were genetically validated by SNP arrays using CytoScan HD (Thermo Fisher Scientific; (GSM888817, GSM888344, GSM888345).

### Macrophage polarization assays

Differentiated M0 macrophages were polarized pharmacologically for 48 h using 1 ng/mL LPS + 20 ng/mL IFN-γ (M1) or 20 ng/mL IL-4 + 20 ng/mL IL-13 (M2). Gene expression of monocytes/macrophages markers (CD68), M1 markers (CD86, HLA-DR) and M2 markers (CD206, CD163) were determined by ddPCR as described below. Data are shown in Supplementary Figure 1.

### PI3K inhibition and Western blot analysis

GBM cells were serum-starved (16 h) and treated with 200 nM wortmannin for 1 h. Untreated or wortmannin-treated control GBM cells were stimulated with insulin (20 nM, 15 min). In extended assays, wortmannin was removed by three washes with PBS and cells were incubated for up to 72 h with fresh media. Proteins were resolved by 12% SDS-PAGE and probed with the following primary antibodies from Cell Signaling Technology: anti-phospho-AKT (Ser473) (Cat#9271; 1:500), total AKT (Cat#9272; 1:1000), cleaved Caspase-3 (Cat#9664; 1:250). GAPDH (1:1000;Cat# sc-365062, RRID:AB_10847862; Santa Cruz Biotechnology) and B-actin (A3854; 1:1000; Sigma-Aldrich) were used as loading controls. HRP-conjugated secondary antibodies included anti-rabbit IgG (A6154; 1:5000; Sigma-Aldrich) and anti-mouse IgG (A9044; 1:5000; Sigma-Aldrich). Bands were visualized with ECL Chemiluminescence Detection Kit (Thermo Fisher Scientific) in an Amersham Imager 600 (GE Healthcare Life Sciences). Chemiluminescent signals were quantified with Sciugo.com.

### Transwell coculture assays

Unpolarized (M0) macrophages were plated in the lower compartment of 6-well transwell plates (1 μm pore size; Corning), while GBM cells were seeded in the upper inserts at a macrophage-to-tumor cell ratio of 2:1. After a 24-h attachment period, cocultures were maintained for three days in medium containing reduced serum levels to allow soluble factor-mediated communication while preventing direct cell contact. For inhibition experiments, GBM cells were exposed to the PI3K inhibitor wortmannin (200 nM) for 1 h prior to coculture, followed by two washes with PBS to remove residual compound. Treated GBM cells were then placed into transwell inserts and cocultured with macrophages under the same conditions. Where indicated, the IL-6 receptor-blocking antibody tocilizumab (500 ng/mL; RoActemra, Roche Farma S.A.) was added to both upper and lower compartments for the duration of the coculture.

### Wound-healing migration assay

Glioblastoma cells were seeded into 48-well plates at a density of 1.2 x 10L cells per well and allowed to form a confluent monolayer overnight. A linear scratch was generated using a sterile 200 μL pipette tip, after which cells were gently rinsed with PBS to remove detached material. Cells were then incubated for 12 h with either standard culture medium, CM derived from 3-day GBM-macrophage cocultures, or CM obtained from cocultures in which GBM cells had been pretreated with wortmannin. Phase-contrast images were acquired at 0, 3, 6, 9, and 12 h using a Leica DMi8 time-lapse microscope. Wound closure was quantified as the percentage reduction in the initial wound area using ImageJ with the Wound_healing_size_tool.ijm plugin, and migration was calculated as: Migration (%) = [(initial wound area - final wound area) / initial wound area] × 100.

### Targeted DNA sequencing and mutation analysis

To identify potential somatic alterations in the RAS-ERK and PI3K-mTOR signaling pathways that may influence autophagy regulation or therapeutic response, we performed targeted DNA sequencing using the Oncomine Comprehensive Assay Plus (Thermo Fisher Scientific). Genomic DNA was extracted from GBM cell lines using the QIAamp DNA Mini Kit (Qiagen) according to the manufacturer’s instructions. Library preparation, target enrichment, and sequencing were carried out using the Ion Torrent Genexus Integrated Sequencer platform following the manufacturer’s protocols.

The Oncomine Comprehensive Assay Plus panel allows detection of mutations, copy number variations (CNVs), and selected gene fusions across >500 cancer-relevant genes, including key components of the RAS-RAF-MEK-ERK and PI3K-AKT-mTOR pathways. Sequence data were aligned to the human reference genome (hg19), and variant calling was performed using the Ion Reporter™ Software (Thermo Fisher Scientific). Variants were annotated and filtered to prioritize pathogenic or likely pathogenic mutations based on ClinVar, COSMIC, and OncoKB databases.

### ddPCR

Transcript expression level was quantified using droplet digital PCR (ddPCR). Reactions were set up in a final volume of 20 μL containing 2x ddPCR Supermix for Probes (no dUTP; Bio-Rad), 20x FAM-labeled TaqMan probe assays targeting genes of interest (CD68, HLA-DR, CD86, CD163, CD206, IL-10, IL-6, MMP9, IDO1, CXCL10, CXCR4, CTLA4, CD274 [PD-L1], CD276 (B7-H3), VEGFA and TGFB1), and a 20x HEX-labeled ACTB probe used as a reference. All reagents were obtained from Bio-Rad. Droplet generation was carried out using the QX200™ Droplet Generator following the manufacturer’s protocol. PCR amplification was performed on a C1000 Touch Thermal Cycler (Bio-Rad), and droplets were subsequently read on a QX200 Droplet Reader. Absolute quantification and data analysis were conducted using QuantaSoft™ software. Gene expression levels are reported as fractional abundance, calculated as the ratio of target transcript copies to ACTB copies. ACTB was selected as the reference gene due to its stable expression across all experimental conditions, including coculture assays.

### mRNA library preparation and sequencing

RNA from GBM cell lines and from M0 macrophages, pharmacologically polarized M2, and M0 cocultured with GBM cells (pre-treated or not with wortmannin) was extracted using the RNeasy Mini Kit (Qiagen) and quantified in Nanodrop. RT-PCR was performed with 100 ng of RNA using the High-Capacity cDNA Reverse Transcription Kit (Applied Biosystems). cDNA was synthesized and stored at 4°C for immediate use or at-80°C for future use.

mRNA libraries were prepared using the Illumina Stranded mRNA Prep kit according to the manufacturer’s protocol. Briefly, mRNAs were purified with oligo-dT beads, fragmented, and hybridized with hexamers to synthesize the first strand of cDNA. Strand specificity was maintained by incorporating dUTPs during second-strand synthesis. Double-stranded cDNA was then ligated to indexed adapters and enriched by PCR. Libraries were quantified by fluorometry, and their size was verified using the Agilent Bioanalyzer. Libraries were pooled and sequenced on a NextSeq 550 platform (Illumina) using 75-cycle single-read sequencing (1x 75 bp).

### mRNAseq analysis

RNA-seq reads were mapped against the human reference genome (GRCh38) with STAR 2.7.8a [25] using ENCODE parameters. Genes were quantified with RSEM 1.3.0 [26] using the gencode 46 annotation. Differential expression was performed with limma [27] using the voom transformation of the counts [28]. Functional enrichment for the up and down-regulated genes was performed separately with g:Profiler [29]. Gene set enrichment analysis was done with the pre-ranked list of genes by the limma t moderated statistic, using FGSEA 1.12.0, using the Reactome pathway database. Macrophage marker genes used for macrophage classification were selected from previously validated literature [30–33] (Supplementary Table 3). For visualization, z-scores were calculated from the mean logCPM values of macrophage M0 and M2 (n = 2), under conditions with or without coculture and treatment (n = 6). Heatmaps were generated using the pheatmap package (v1.0.13).

### Statistical analysis

Data was generated from three biological independent experiments. Results are presented as mean ± standard error of the mean (SEM). For two-group comparisons, an unpaired two-tailed t-test was used. Statistical significance among three or more groups was assessed using one-way analysis of variance (ANOVA) followed by Tukey’s post hoc test. Statistical significance was set at p ≤ 0.05, with significance levels denoted as follows: *p < 0.05; **p < 0.01; ***p < 0.001; ****p < 0.0001. Graphs were visualized using R, R2 or GraphPad. The datasets are publicly available under “GBM (Wortmannin tr) - FontdeMora - 24” and “GBM (Macrophage tr) - FontdeMora - 16”. Single cell analyses were performed in R 4.5.2.

## RESULTS

### MRI-defined vascular phenotypes reveal macrophage enrichment and PI3K-and cytokine-related signaling in primary, IDH-wild type GBM

To investigate whether vascular heterogeneity reflects distinct immune and molecular features in GBM, we stratified 15 IDH-wild type GBM patients from TCGA-GBM database using MRI-derived vascular biomarkers obtained through the ONCOhabitats platform. This automated radiomic approach integrates multi-parametric MRI data to quantify intratumoral heterogeneity and objectively calculate relative rCBV, providing a reproducible and operator-independent assessment of vascularity. ONCOhabitats enabled objective quantification of tumor vascularity for integration with transcriptomic analyses. Based on the rCBV median threshold at the HAT [16, 17], tumors were stratified as HV (rCBV>=5.9, *n* = 8) or MV (rCBV<5.9, *n* = 7) (Figure 1A). As expected, Kaplan-Meier analysis revealed that patients with HV tumors exhibited shorter overall survival than those with MV tumors (Figure 1B).

**Figure 1.**
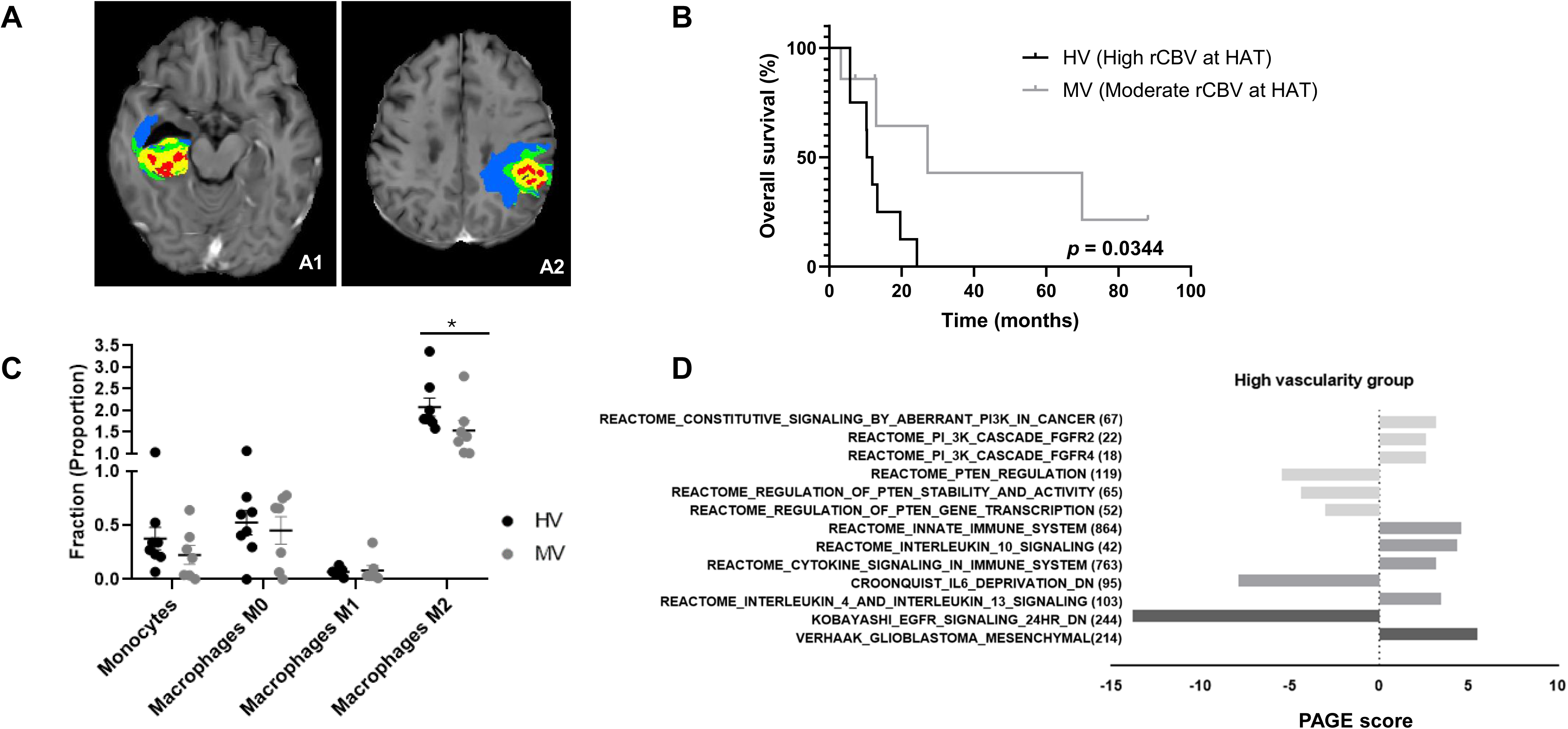
**Highly vascular GBMs display increased macrophage infiltration and transcriptional enrichment of PI3K and immune signaling pathways**. (A) Axial T1-weighted contrast-enhanced (T1c) MR images showing GBM tumor delineation using ONCOhabitats in patients with high (A1) and moderate (A2) vascularity in the HAT habitat (in red). (B) Kaplan-Meier overall survival analysis of glioblastoma patients stratified by vascularity. The HV group includes patients with rCBV HAT ≥ 5.9 (n = 8), whereas the MV group includes patients with rCBV HAT < 5.9 (n = 7). Survival curves were compared using the log-rank (Mantel-Cox) test (χ² = 4.475, *p* = 0.0344). The HV group exhibited a significantly higher risk of death compared with the MV group (hazard ratio = 3.95; 95% CI, 1.11-14.13). (C) CIBERSORTx deconvolution of bulk RNA-seq data showing the relative abundance of macrophage populations between HV and MV groups (unpaired two-tailed t-test, *p* = 0.0289). (D) Parametric Analysis of Gene Set Enrichment (PAGE) performed in R2 software, highlighting transcriptional programs enriched in HV tumors, including PI3K/EGFR activation, PTEN downregulation, and cytokine-related signaling pathways (IL-10, IL-13, and innate immune system).

To assess immune cell composition, CIBERSORTx deconvolution of bulk RNA-seq data revealed a significantly higher abundance of macrophages in HV compared with MV tumors (unpaired two-tailed t-test, *p* = 0.0289; Figure 1C), suggesting that vascular density associates with increased myeloid infiltration. Subsequent transcriptomic profiling using PAGE analysis in R2 software identified enrichment in HV tumors of signaling pathways related to PI3K and EGFR activation, loss of *PTEN* regulation, and cytokine-mediated processes including IL-10, IL-13, and innate immune responses (Figure 1D, see also Supplementary Figure 2). Together, these data indicate that highly vascular GBMs are characterized by enhanced macrophage presence and upregulation of PI3K-and cytokine-associated transcriptional programs.

### Mesenchymal GBM subtype associates with M2 macrophage enrichment and PI3K pathway alterations

To extend the observations from our 15-patient cohort, we analyzed 147 IDH-wild-type GBMs from TCGA, of which 130 contained RNA-sequencing data and were included in transcriptomic analyses. Patients were classified into the Verhaak molecular subtypes, mesenchymal (MES), classical (CL), neural (NE) and proneural (PN), using the corresponding Bioconductor pipeline (17 cases lacked RNA-seq data and were excluded from the analysis). We next applied CIBERSORTx to estimate immune cell composition across subtypes. The MES subtype exhibited markedly higher infiltration of M0 and M2 macrophages, as well as an overall increase in total monocyte content, compared with CL and PN tumors (Figure 2A). In addition, analysis of PI3K pathway alterations revealed that *PIK3CA* mutations and *PTEN* loss were significantly enriched in MES GBMs (Figure 2B). The classification of all cases according to the original Verhaak scheme is shown in Figure 2C, and the code used for transcriptomic subtype assignment is provided in the Materials and Methods section. These findings indicate that the mesenchymal transcriptional program in GBM is linked to macrophage polarization and activation of the PI3K signaling pathway.

**Figure 2.**
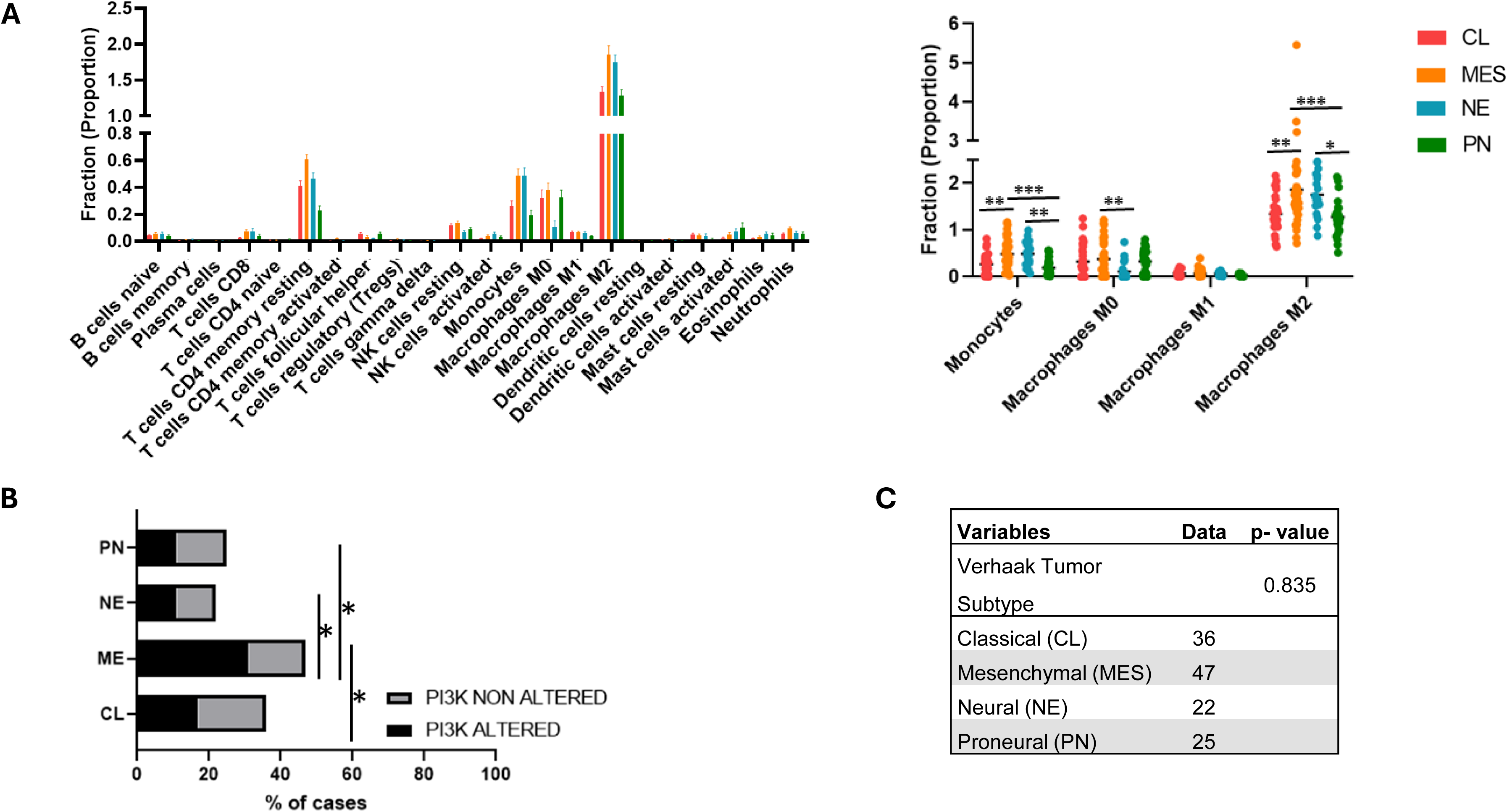
Mesenchymal GBM subtype is associated with M2 macrophage enrichment and PI3K pathway alterations. (A) Estimation of immune cell populations across GBM molecular subtypes in the TCGA cohort using CIBERSORTx (*n* = 130 IDH-wild-type cases). The mesenchymal (MES) subtype displays higher infiltration of M0 and M2 macrophages and increased total monocyte abundance compared with proneural (PN), neural (NE) and classical (CL) subtypes. (B) Distribution of PI3K pathway alterations (including *PIK3CA* mutation or *PTEN* loss) across GBM molecular subtypes, showing enrichment in MES tumors. (C) Classification of GBM samples from the GBM cohort according to the original Verhaak transcriptomic subtypes using the Bioconductor pipeline. Data are represented as mean ± SEM. Statistical significance was determined by one-way ANOVA with multiple comparisons (*p* ≤ 0.05; \**p* < 0.01; \*\**p* < 0.001; \*\*\**p* < 0.0001).

### Molecular profiling of GBM cell lines confirms distinct PI3K pathway alterations

To characterize baseline differences in PI3K pathway status among glioblastoma models displaying mesenchymal-like features, we performed targeted DNA sequencing using the Oncomine Comprehensive Assay Plus panel. Analysis confirmed that U87 harbors a *PTEN* splice variant (NM_000314.8) with 100 % allele frequency, consistent with its known loss of PTEN function and constitutive PI3K/AKT activation. LN-18 cells carried a *PIK3CB* p.Glu1051Lys mutation (allele frequency 100 %), while LN-229 cells were wild type for genes within the PI3K pathway. These findings corroborate previous reports and justify their use as complementary models exhibiting different baseline PI3K activity states. Raw variant data are provided in Supplementary File 1.

### Validation of macrophage polarization and PI3K inhibition kinetics prior to coculture assays

Before establishing macrophage-GBM cocultures, pharmacological differentiation of THP-1 monocytes into macrophages and their subsequent polarization toward M1 or M2 phenotypes was confirmed (Supplementary Figure 3). The PI3K inhibition conditions were then optimized in three GBM cell lines using the irreversible inhibitor wortmannin to determine the persistence of pathway blockade after short-term exposure. A 1-h treatment followed by drug removal effectively suppressed AKT phosphorylation upon insulin stimulation and maintained reduced p-AKT levels for up to 72 h, indicating a prolonged inhibitory effect on PI3K signaling (Supplementary Figure 4A-B). Cleaved caspase-3 analysis confirmed that these conditions did not induce marked apoptosis (Supplementary Figure 4C). These results defined the optimal parameters for subsequent macrophage-GBM coculture experiments.

### Independent single-cell transcriptomic analysis identifies inflammatory programs associated with MES-like malignant glioblastoma cells

To determine whether the inflammatory programs identified in our experimental models were conserved in human disease, we interrogated the independent Smart-seq2 single-cell RNA-sequencing dataset from Neftel *et al*. [23]. After removal of non-malignant cells (macrophages, T cells and oligodendrocytes) according to the Neftel classification, 6,860 malignant cells from 28 primary glioblastomas were retained (Figure 3A; full cell-type classification of all 7,930 cells in Supplementary Figure 5A). Within these cells, the mesenchymal (MES-like) module score and the tumor-macrophage communication signature score, were concentrated in overlapping regions of the malignant-cell UMAP (Figure 3B,C). Within individual tumors, MES-like and tumor-macrophage communication signature scores were positively correlated in the majority of patients (median within-tumor Spearman ρ = 0.28), and a linear mixed model with a by-patient random slope and intercept confirmed a highly significant positive fixed effect of the MES-like score (p = 1.6 × 10⁻¹¹; Figure 3D), indicating that the association is not driven by between-patient differences. The MES-like transcriptional program and tumor-macrophage communication signatures contained no overlapping genes, excluding gene-set overlap as an explanation for the observed association. Across tumors, the proportion of MES-like malignant cells (four-state argmax) was strongly correlated with the mean tumor-macrophage communication signature score (Spearman ρ = 0.82, p = 8.0 × 10⁻L; Figure 3E). The same relationship held for an NF-κB inflammatory program included as an orthogonal robustness check: the NF-κB score co-localized with the MES-like program on the malignant-cell UMAP (Supplementary Figure 5B) and was positively associated with it both within tumors (median Spearman ρ = 0.24; linear mixed model p = 3.0 × 10⁻¹¹; Supplementary Figure 5C) and at the tumor level (Spearman ρ = 0.80, p = 2.7 × 10⁻L; Supplementary Figure 5D). Analysis of the remaining malignant transcriptional states showed that tumors enriched for the astrocyte-like (AC-like) state exhibited weaker positive correlations with the mean NF-κB and tumor-macrophage communication signature scores (Spearman ρ = 0.59 and 0.62, respectively) compared to the MES-like score, whereas tumors enriched for the neurodevelopmental OPC-like and NPC-like states showed the opposite pattern (OPC: ρ = −0.77 and −0.76; NPC: ρ = −0.57 and −0.57; Supplementary Figure 5E-G, Supplementary Table 2); as the four state fractions are compositionally constrained to sum to 100%, these non-MES associations should be read as relative rather than independent effects, but their concordant direction across both the inflammatory and the communication readout supports a specific link between the MES-like axis and inflammatory tumor-macrophage signaling.

**Figure 3.**
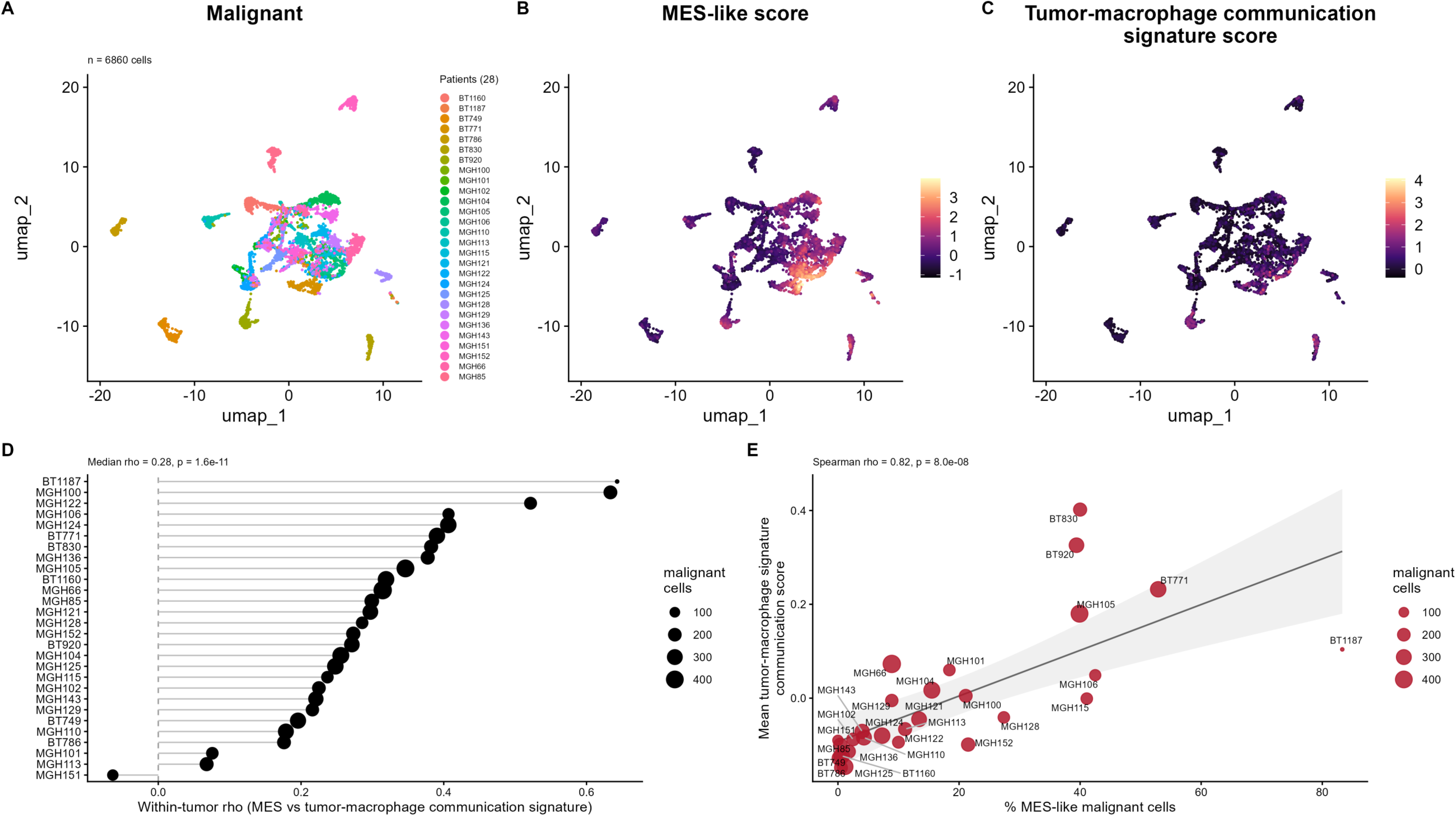
Malignant GBM cells with a stronger mesenchymal program show higher tumor-macrophage communication signature activity in single-cell cohort. (A) UMAP representing the 6,860 malignant cells retained across 28 primary glioblastomas (GEO GSE131928, Smart-seq2 IDH-wildtype cohort) after removal of non-malignant lineages, colored by patient of origin. (B) UMAP representing malignant cells colored by the mesenchymal (MES-like) module score. (C) UMAP representing malignant cells colored by the tumor-macrophage communication signature score. (D) Within-tumor Spearman correlations (ρ) between the MES-like score and the communication score, one point per patient; point size denotes the number of malignant cells per tumor. The subtitle reports the median within-tumor ρ and the fixed-effect p-value from a linear mixed model with a by-patient random slope and intercept. (E) Tumor-level association between the percentage of MES-like malignant cells (Neftel four-state argmax) and the mean communication score; one point per tumor, point size denotes malignant-cell number, line and shaded band show the linear fit with 95% confidence interval, and the subtitle reports the Spearman ρ and p-value (n = 28 tumors).

Overall, these independent single-cell analyses demonstrate that inflammatory transcriptional programs are selectively associated with MES-like malignant glioblastoma cells at both the single-cell and patient levels.

### GBM-macrophage crosstalk induces macrophage polarization through PI3K-dependent signaling

To explore how GBM cells modulate macrophage polarization, we cocultured unpolarized M0 macrophages with three GBM cell lines (U87, LN-18, and LN-229) using a transwell insert-based system that prevents physical contact but allows soluble factors to diffuse between compartments (Figure 4A). After three days, gene expression was analyzed by ddPCR to assess cytokine induction and polarization markers. Coculture markedly increased IL-6 expression in both macrophages and GBM cells, indicating a strong bidirectional cytokine response (Figure 4B,C). Although IL-6 is traditionally considered a pro-inflammatory mediator, in tumor contexts, including GBM, it functions predominantly as an immunosuppressive cytokine promoting M2-like macrophage polarization [34]. Notably, pretreatment of GBM cells with wortmannin attenuated IL-6 induction in tumor cells and reduced IL-6 levels in macrophages, suggesting that tumor-intrinsic PI3K signaling contributes to the establishment of this IL-6-rich microenvironment. IL-10 expression was strongly induced in macrophages and to a lesser extent in GBM cells. Wortmannin treatment reduced IL-10 expression in macrophages cocultured with U87 and LN-18 cells but not in those with LN-229 cells, highlighting tumor line-specific differences in PI3K pathway regulation.

**Figure 4.**
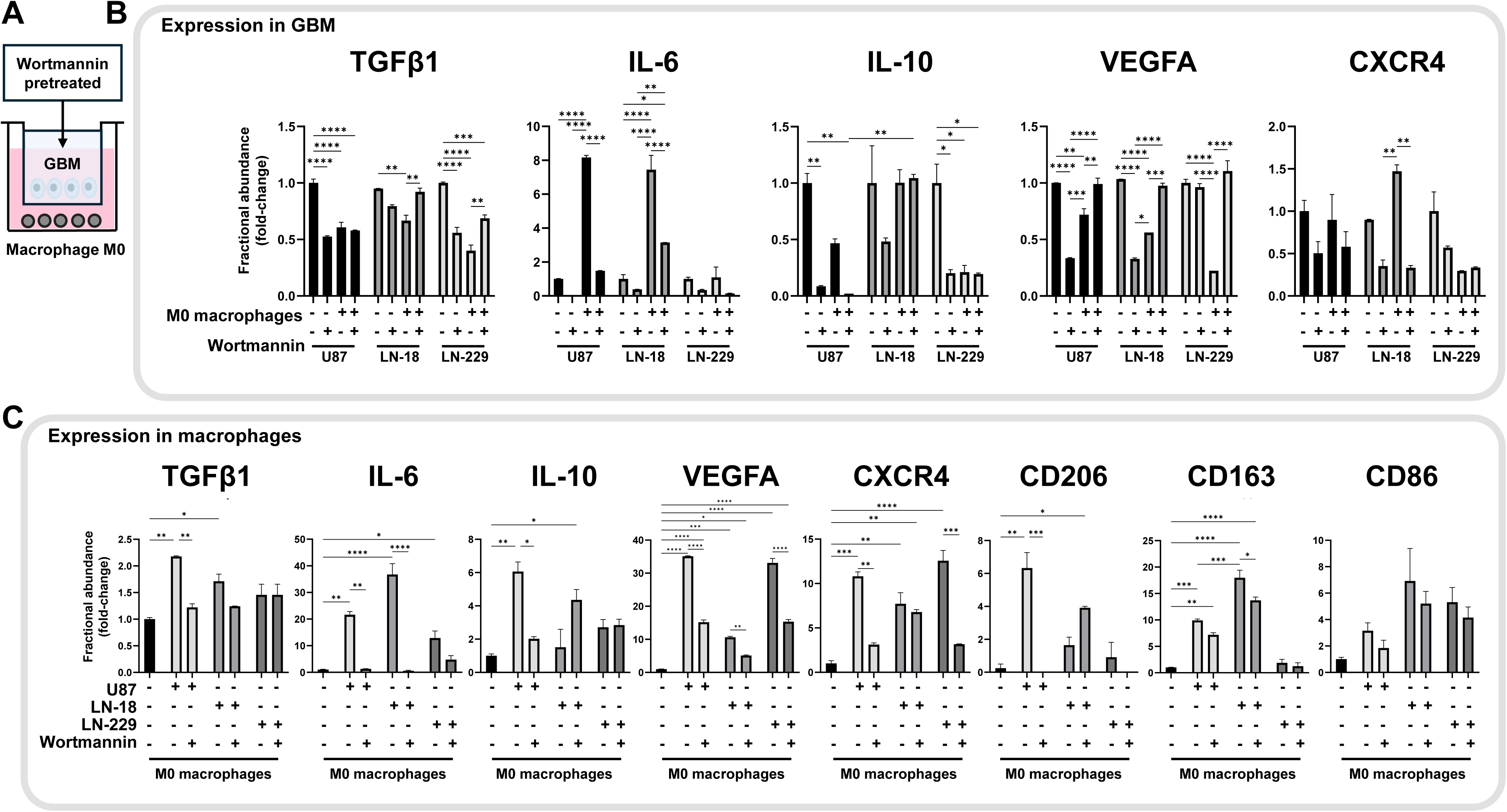
Coculture of M0 macrophages with glioblastoma cell lines induces polarization toward M2-like phenotypes. (A) Schematic of the insert-based coculture system, where M0 macrophages (unpolarized, THP-1-differentiated as described in Supplementary Figure 1) are cocultured with glioblastoma cell lines in separate chambers of a transwell insert (1 µm pore size), preventing physical contact but allowing soluble factors to exchange between compartments. Arrows indicate the workflow and downstream analyses performed on cells from each side of the insert. (B) Expression of TGFB1, IL-6, IL-10, VEGFA and CXCR4 was quantified by ddPCR under control conditions or following wortmannin treatment (200 nM, 1 h). After treatment, tumor cells were washed three times with fresh medium and subsequently cultured with or without macrophages, as indicated. (C) Cytokine and polarization marker expression in macrophages measured by ddPCR after 3 days of coculture. Expression TGFB1, IL-6, IL-10, VEGFA and CXCR4, as well as M1-associated (CD86) and M2-associated (CD206, CD163) markers, was evaluated. Data are represented as fractional abundance relative to the housekeeping gene β-actin. Fold change was calculated as the ratio between expression in the coculture condition and their respective non-cocultured controls (glioblastoma cells or M0 macrophages). Values represent means ± SEM of duplicates from up to four independent experiments. *p* ≤ 0.05; \**p* < 0.01; \*\**p* < 0.001; \*\*\**p* < 0.0001 (one-way ANOVA).

At the phenotypic level, macrophages cocultured with GBM cells exhibited increased expression of CD163 and CD206, consistent with an M2-like polarization pattern, while CD86 expression was also elevated, suggesting a partially activated or “hybrid” phenotype (Figure 4C). Importantly, inhibition of PI3K signaling modulated the expression of macrophage polarization markers, with the most pronounced effects observed in U87 cells and more variable responses in LN18 and LN229 models. Collectively, these data support a model in which tumor-intrinsic PI3K signaling promotes the production of IL-6 and IL-10, thereby contributing to an immunosuppressive, M2-like macrophage phenotype and highlighting PI3K-dependent cytokine signaling as a key determinant of the GBM microenvironment.

### PI3K inhibition in GBM cells reduces the expression of immune checkpoint molecules in tumor-associated macrophages

Having established that PI3K signaling modulates the reciprocal cytokine exchange between GBM cells and macrophages (Figure 4), we next sought to determine whether this pathway also governs the expression of immune checkpoint molecules that sustain macrophage-mediated immunosuppression. Since cytokines such as IL-6 and IL-10 are known regulators of checkpoint expression, we examined whether PI3K inhibition in GBM cells could disrupt this signaling axis and consequently alter the tumor-associated macrophage phenotype. To this end, we measured the expression of PD-L1 (CD274), CTLA-4, IDO1, and CD276 (B7-H3) in the coculture system with wortmannin pretreated or untreated GBM cells (Figure 5).

**Figure 5.**
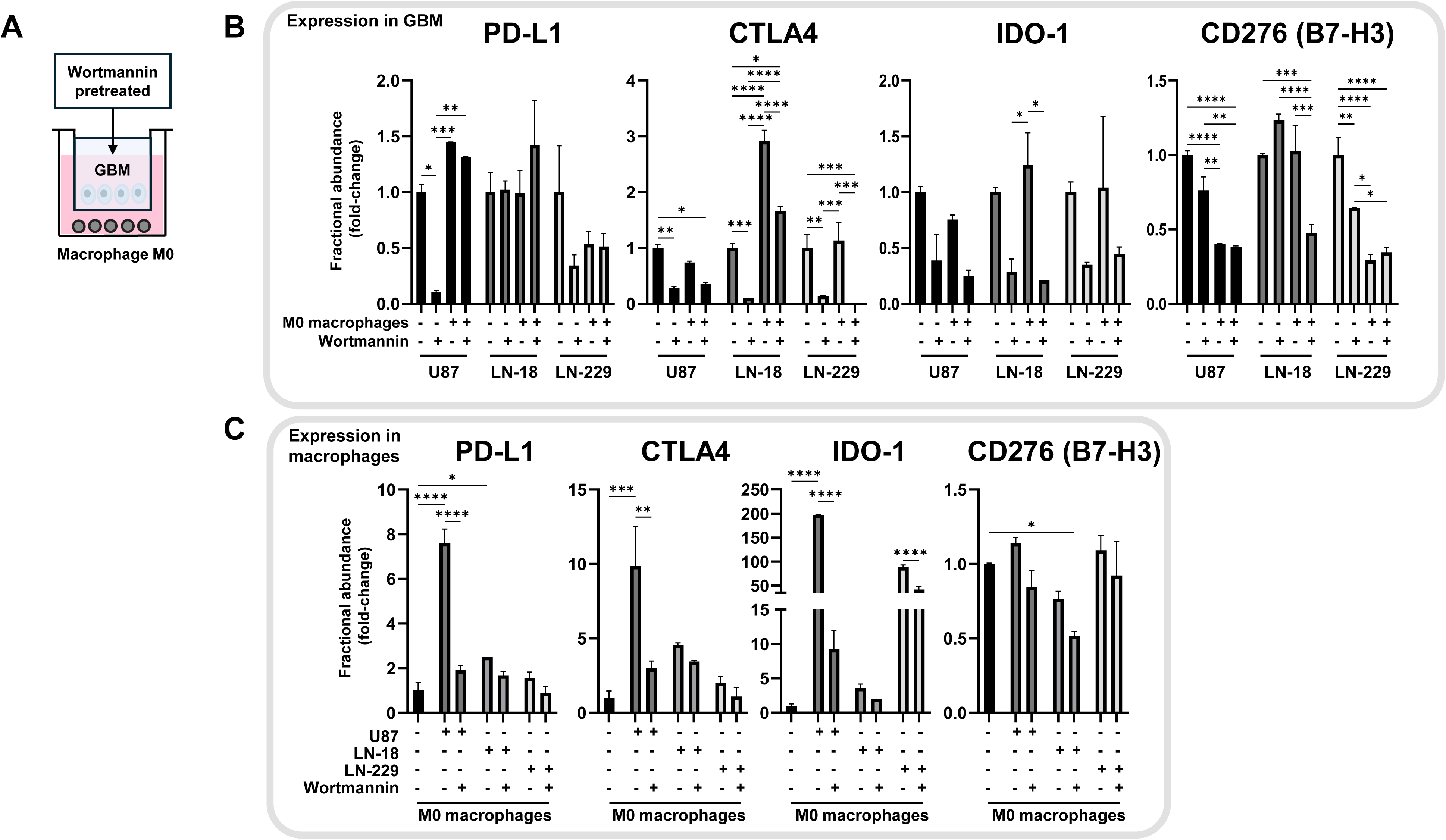
**PI3K inhibition in GBM cells diminishes the induction of immune checkpoint genes in both tumor cells and macrophages**. (A) Schematic representation of the transwell coculture assay. M0 macrophages (unpolarized, THP-1-derived) were cultured with GBM cells in separate chambers of a 1 µm-pore insert, preventing physical contact while allowing exchange of soluble factors. GBM cells were pre-treated with wortmannin (200 nM, 1 h) to inhibit PI3K signaling. (B) Expression of PD-L1 (CD274), CTLA-4, IDO1, and CD276 (B7-H3) in GBM cells before and after coculture with M0 macrophages. Tumor cells were treated with wortmannin (200 nM, 1 h) or vehicle control, washed three times with fresh medium, and subsequently cocultured with or without macrophages. (C) Expression of the same markers in macrophages after three days of coculture with GBM cells pre-treated or not with wortmannin. Data are represented as fractional abundance relative to the housekeeping gene β-actin. Fold change was calculated as the ratio between expression in cocultured and non-cocultured controls (GBM cells or M0 macrophages). Values represent means ± SEM of duplicates from three independent experiments. *p* ≤ 0.05; \**p* < 0.01; \*\**p* < 0.001; \*\*\**p* < 0.0001 (one-way ANOVA).

Using the same transwell coculture system described above, M0 macrophages were cultured with GBM cells for three days (Figure 5A), and gene expression was quantified by ddPCR (Figure 5B,C). Macrophages exposed to tumor cell lines exhibited a significant increase in the expression of the immunoregulatory markers PD-L1 (CD274), IDO1, and CTLA-4, whereas CD276 (B7-H3) levels remained largely unchanged (Figure 5C). Among the GBM lines, the strongest induction of macrophage checkpoint genes occurred in cocultures with U87 cells (Figure 5B), which harbor a PTEN mutation and display constitutive PI3K-dependent AKT activation, as well as a sustained inhibition by wortmannin (Supplementary Figure 4). PI3K inhibition in GBM cells reduced the induction of selected immune checkpoint markers, including PD-L1 and IDO1, with more variable effects observed for CTLA-4 and across different GBM models.

Conversely, GBM cells exposed to macrophages showed more modest transcriptional changes than macrophages themselves, although wortmannin treatment significantly reduced checkpoint expression in all cell lines. CD276 (B7-H3) expression decreased in U87 and LN-229 cells following macrophage exposure but was not further reduced by wortmannin, suggesting a limited contribution of PI3K signaling to CD276 regulation.

Collectively, these findings indicate that PI3K activity in GBM cells contributes to the regulation of macrophage expression of immune checkpoint molecules, highlighting its pivotal role in driving immunosuppressive communication within the GBM microenvironment.

### Tumor-macrophage crosstalk enhances GBM cell migration through PI3K-dependent signaling

To determine whether the interaction between GBM cells and macrophages generates soluble factors that modulate tumor aggressiveness, we assessed the migration capacity of naïve GBM cell lines exposed to CM derived from the three-day cocultures with GBM cells, either pretreated or not with wortmannin.

A wound-healing assay performed over 12 h revealed that CM from GBM-macrophage cocultures markedly increased the migration rate of all GBM lines, indicating that the crosstalk between these two cell types releases soluble factors that promote tumor motility (Supplementary Figure 6). In contrast, CM obtained from cocultures in which tumor cells had been pre-treated with wortmannin failed to stimulate migration, showing rates comparable to control media (Supplementary Figure 6). These results demonstrate that PI3K signaling in GBM cells contributes to the secretion of pro-migratory factors during macrophage interaction, linking oncogenic signaling to microenvironment-mediated tumor dynamics.

As an internal control, direct treatment of GBM cells with wortmannin (without CM) also reduced migration (Supplementary Figure 6), consistent with the role of PI3K/AKT signaling in regulating intrinsic motility. Together, these findings indicate that PI3K activation drives both the release and responsiveness to paracrine pro-migratory cues, establishing a positive feedback loop that reinforces GBM aggressiveness.

### Tumor-intrinsic PI3K activity drives IL-6-dependent paracrine signaling that shapes immunosuppressive macrophage states

To investigate how tumor-intrinsic PI3K activity shapes macrophage phenotype and paracrine signaling, we performed two complementary sets of experiments. First, GBM cell lines (U87, LN-18, LN-229) were pretreated or not with wortmannin, washed twice to remove residual inhibitor, and then placed in transwell coculture with M0 macrophages (macrophages were never directly exposed to wortmannin). Macrophage lysates were analyzed by immunoblot and RNA-seq (biological duplicates) to assess signaling and global transcriptional responses (Figure 6).

**Figure 6.**
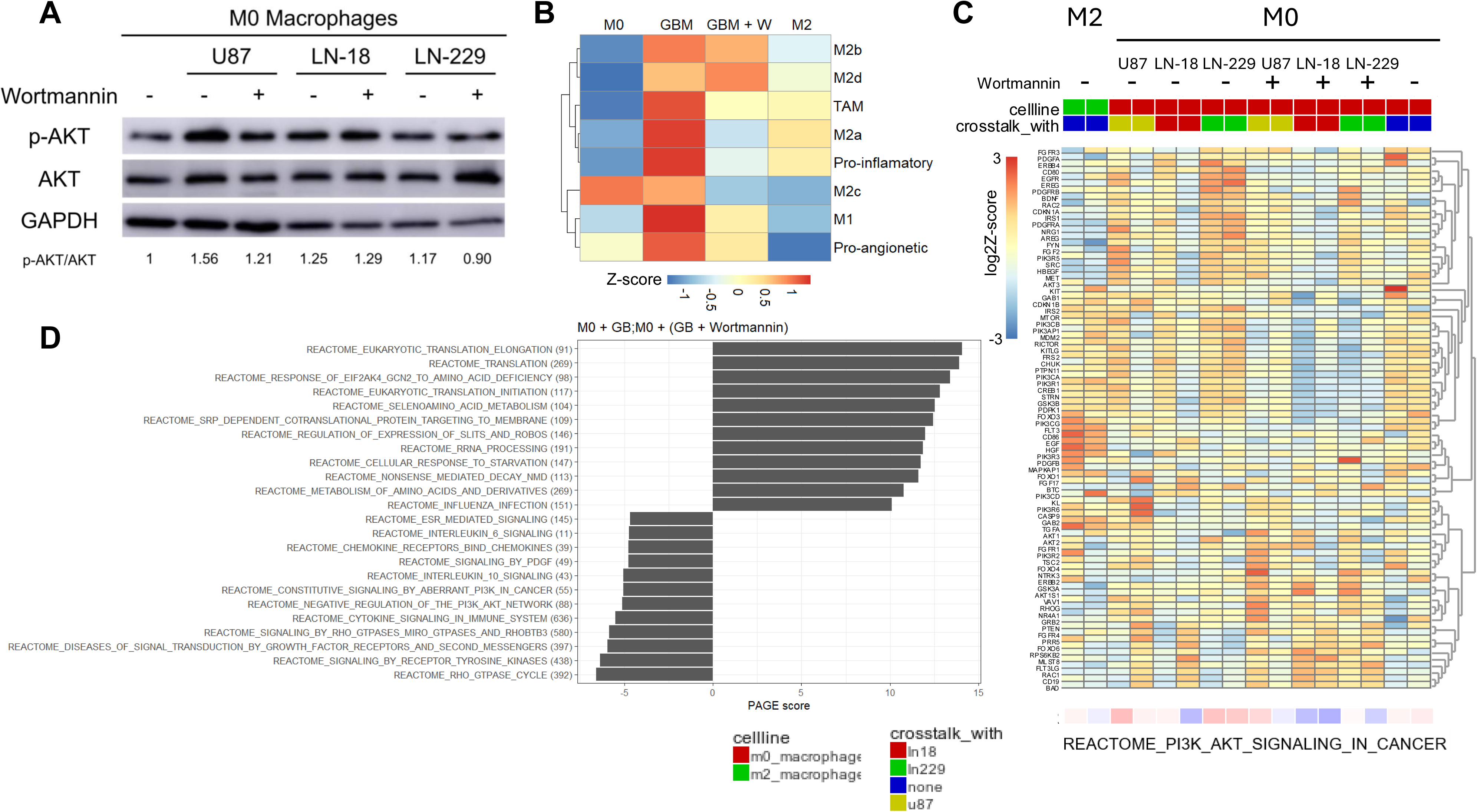
Tumor-intrinsic PI3K activity modulates macrophage transcriptional programs with limited effect on macrophage AKT phosphorylation. (A) Western blot analysis of phospho-AKT (p-AKT Ser473), total AKT and GAPDH in macrophages after 3-day transwell coculture with U87, LN18, or LN229 GBM cells that were pre-treated or not with wortmannin (200 nM, 1 h) and washed twice prior to coculture with macrophages. Inhibition of AKT phosphorylation in GBM cells was highly efficient and sustained in time (Supplementary Figure 4). Representative blots of macrophages’ lysates and quantification of p-AKT/total AKT/GAPDH ratios are shown; changes in macrophage p-AKT upon coculture are subtle and variable across lines. Prior tumor-cell wortmannin treatment produces only minor attenuation of macrophage p-AKT. (B) Heatmap of macrophage transcriptomic responses (RNA-seq, biological duplicates for each cell line and condition) to coculture with glioblastoma cells pre-treated or not with wortmannin. Marker genes were selected as described in Methods and are listed in Supplementary Table 3. For visualization, Z-scores were calculated from the mean logCPM values of M0 and pharmacologically polarized M2 macrophages (n = 2) across conditions with or without coculture and tumor-cell wortmannin treatment (n = 6). Rows are marker genes for macrophage subtypes; columns are macrophage samples/conditions. Color bar indicates Z-score scale. (C) Transcriptomic profiling (RNA-seq, biological duplicates) of macrophages: M0, pharmacologically polarized M2, and M0 macrophages cocultured with each GBM line (pre-treated or not with wortmannin). Heatmap shows Reactome “PI3K-AKT signaling in cancer” gene set Z-scores (R2 Genomics dataset: *GBM (Wortmannin tr) - FontdeMora - 24*). Macrophages exposed to GBM cells with inhibited PI3K display reduced Z-scores for PI3K-AKT-related transcriptional programs (color bar indicates Z-score scale). Data support that tumor-cell PI3K activity modulates macrophage PI3K/AKT-related transcriptional output despite only modest changes in steady-state p-AKT levels detected by immunoblot. (D) Parametric Analysis of Gene-set Enrichment (PAGE) analysis of macrophage transcriptional programs following coculture with GBM cells pre-treated or not with wortmannin. PAGE was performed using the R2 Genomics platform to compare macrophages cocultured with each GBM cell line under control conditions versus macrophages exposed to GBM cells pre-treated with wortmannin. Top 12 up and down regulated REACTOME gene sets are shown. Pathways associated with negative regulation of PI3K signaling, as well as IL-6 signaling, IL-10 signaling, and cytokine signaling in the immune system, were diminished under wortmannin conditions. These results indicate that PI3K activity in GBM cells drives the induction of macrophage transcriptional programs linked to translation, cytokine responsiveness, and PI3K/AKT-associated regulatory networks.

By immunoblot, macrophage p-AKT showed only subtle and variable changes across coculture conditions, and prior tumor-cell wortmannin exposure produced at best a minor attenuation of p-AKT (Figure 6A). To obtain a broader view of macrophage phenotypic shifts, we next evaluated curated macrophage gene signatures derived from established M0 and M2 polarization markers. Heatmap visualization of these signatures demonstrated that coculture with PI3K-competent tumor cells consistently increased expression of M2-associated genes, whereas macrophages exposed to wortmannin-pretreated tumor cells displayed attenuated induction across multiple marker sets (Figure 6B).

Transcriptomic profiling revealed a robust effect of tumor PI3K on macrophage programs: macrophages cocultured with PI3K-competent tumor cells displayed elevated Z-scores for the “PI3K-AKT signaling in cancer” Reactome gene set, whereas those exposed to wortmannin-pretreated tumor cells showed reduced Z-scores (Figure 6C). PAGE analysis comparing macrophages exposed to untreated GBM cells (M0+GB) versus wortmannin-pretreated GBM cells (M0+(GB+Wortmannin)) identified strong enrichment, when tumor PI3K was intact, of cytokine-driven pathways (IL-6, IL-10, IL-4/IL-13, IL-1, IFN-γ) as well as anabolic and translational pathways (Figure 6D; Supplementary Figure 7). These transcriptional signatures reflect a mixed but predominantly M2-like TAM phenotype, consistent with the concurrent elevation of CD163, CD206, IL-6, and IL-10 detected by ddPCR (Figure 4) and align with established M2a, M2c, and M2d polarization programs [35, 36]. The coexistence of IL-4/IL-13-, IL-10-, and IL-6-associated signatures is consistent with the known hybrid states of tumor-associated macrophages *in vivo* [37]. IL-10 signaling dropped from the top-ranked pathway to outside the leading gene sets, and IL-4/IL-13, IFN-γ, and IL-1 signatures showed substantial decreases in PAGE scores (Supplementary Figure 7). No cytokine-driven Reactome pathway was enriched when comparing macrophages exposed to wortmannin-pretreated tumor cells versus unpolarized M0 controls. Together, these data indicate that tumor-intrinsic PI3K signaling contributes to the induction of M2-like macrophage transcriptional programs.

Second, to investigate whether IL-6, that was strongly induced in both compartments (Figures 4,5) and its associated Reactome signatures were suppressed by PI3K inhibition, mediates the downstream paracrine signaling and functional effects of tumor-macrophage crosstalk, we generated CM from cocultures (Figure 7A) and used this CM to treat naïve GBM cells in the presence or absence of tocilizumab (IL-6 receptor inhibitor). CM from cocultures consistently increased AKT phosphorylation in recipient tumor cells (Figure 7A). By contrast, treatment of GBM cells with tocilizumab largely abolished CM-driven AKT phosphorylation, supporting a prominent role for IL-6 as a component of the conditioned-media-driven PI3K/AKT response (Figure 7A). Notably, LN-229 cells (PTEN wild type, no PI3K mutation) showed minimal AKT activation upon tocilizumab treatment, supporting a predominant role for IL-6-driven paracrine signaling in cell lines with otherwise low intrinsic PI3K activity.

**Figure 7.**
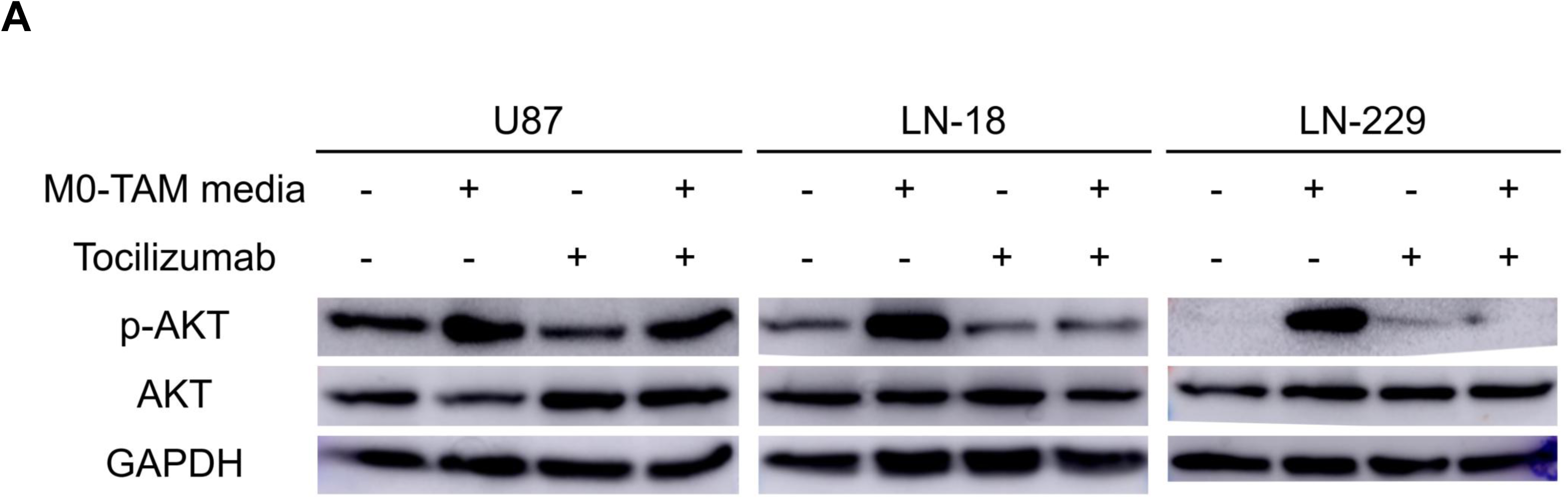
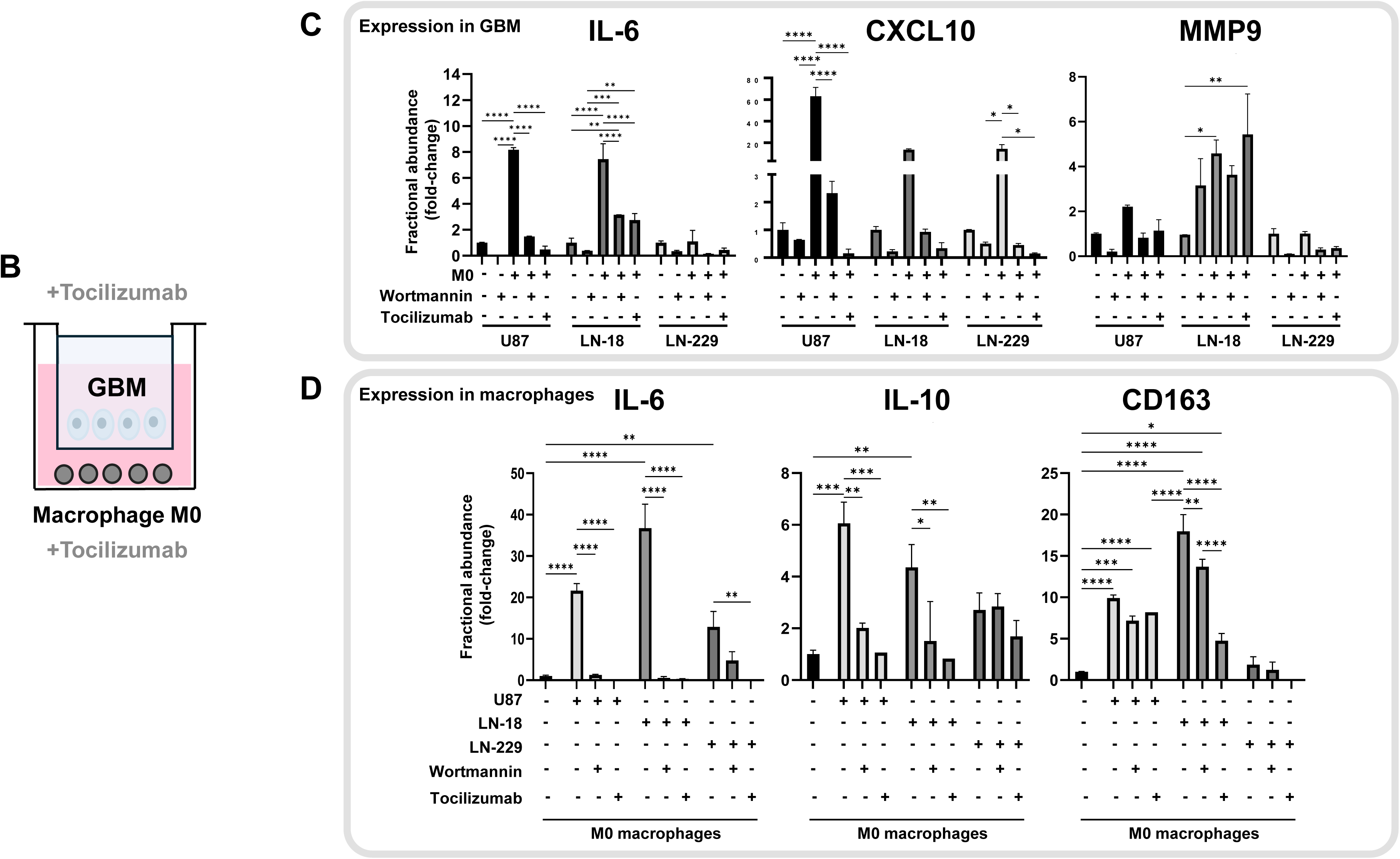
IL-6 blockade abrogates conditioned-media activation of AKT and reduces cytokine and polarization signatures in glioblastoma-macrophage co-cultures. (A) Western blot of phospho-AKT (p-AKT) and total AKT in glioblastoma cells treated with conditioned media (CM) from macrophage or tumor cultures; naïve GBM cells were untreated or treated with 500 ng/ml tocilizumab. Representative blots are shown. (B) Experimental schematic of transwell coculture with M0 macrophages and glioblastoma cells treated or not with tocilizumab. (C) Expression of IL-6, CXCL10, and MMP9 in glioblastoma cells, measured by ddPCR, after three days of co-culture ± tocilizumab (500 ng/ml). (D) Expression of IL-6, IL-10, and CD163 in macrophages, measured by ddPCR, after three days of co-culture ± tocilizumab (500 ng/ml). Data are represented as fractional abundance relative to β-actin; fold change was calculated relative to non-co-cultured controls. Values represent means ± SEM of duplicates from up to four independent experiments. *p* ≤ 0.05; \**p* < 0.01; \*\**p* < 0.001; \*\*\**p* < 0.0001 (one-way ANOVA).

Finally, we tested the impact of IL-6 blockade during transwell coculture. Tocilizumab significantly reduced IL-6 and CXCL10 expression in tumor cells and lowered MMP9 in most lines (except LN-18, suggesting some IL-6-independent induction in this cell line) while macrophage expression of IL-6, IL-10, and the M2 marker CD163 was markedly diminished (Figure 7C,D). Together with the PAGE/RNA-seq data (Figure 6), these results indicate that tumor-intrinsic PI3K activity drives generation of an IL-6-rich secretome that both (i) activates PI3K/AKT signaling in tumor cells in a paracrine manner and (ii) reprograms macrophages toward an M2-like, immunosuppressive transcriptional state. Importantly, CM from cocultures derived from wortmannin-pretreated tumor cells shows reduced ability to stimulate AKT and cytokine programs in macrophages, underscoring that tumor PI3K activity plays an important role in generating these functional paracrine cues.

## DISCUSSION

In this study, we demonstrate that PI3K activation promotes reciprocal tumor-macrophage crosstalk associated with inflammatory signaling, vascular remodeling and MES-like malignant-cell states in glioblastoma. Using complementary *in vitro*, radiogenomic, transcriptomic and single-cell approaches, we identified macrophage-derived inflammatory signaling as a major consequence of PI3K activation. Importantly, analysis of an independent single-cell glioblastoma cohort demonstrated that malignant MES-like cells are consistently associated with enhanced NF-κB activation and tumor-macrophage communication programs, supporting the clinical relevance of our experimental findings.

Although PI3K signaling has traditionally been viewed as a tumor cell-autonomous pathway controlling proliferation, metabolism and survival, our data support an additional role in shaping the inflammatory tumor microenvironment. Rather than acting exclusively within malignant cells, PI3K activation promoted macrophage polarization toward an immunosuppressive phenotype characterized by increased cytokine production and enhanced capacity to induce MES-like transcriptional reprogramming in neighboring tumor cells.

The independent single-cell analyses demonstrate that inflammatory signaling is not uniformly distributed across malignant glioblastoma cells but is preferentially associated with the MES transcriptional state. Importantly, these associations were observed both across individual malignant cells within tumors and at the patient level, supporting the robustness of the inflammatory-MES relationship across different levels of biological organization. Importantly, the MES-like transcriptional program and tumor-macrophage communication signatures were constructed from non-overlapping gene sets, minimizing the possibility that the observed associations reflect gene-set redundancy rather than biological relationships. While these analyses do not establish causality, they are consistent with our functional observations and support a model in which inflammatory tumor-macrophage crosstalk contributes to the maintenance of MES-like malignant-cell states. Importantly, analyses across all four malignant transcriptional states revealed that inflammatory signaling was not a general feature of glioblastoma cells but preferentially associated with the MES program, whereas OPC-like and NPC-like states showed the opposite trend. This observation supports the specificity of the inflammatory circuitry identified in our experimental models.

Mechanistically, PI3K activity in glioblastoma cells promoted macrophage polarization through a cytokine-dependent paracrine program involving IL-6 and IL-10. Tumor-intrinsic PI3K activity was associated with increased expression of these cytokines, induction of macrophage immune checkpoint markers, and transcriptional programs enriched for cytokine signaling and PI3K/AKT-related pathways. Conversely, transient pharmacologic inhibition of PI3K in tumor cells, or blockade of IL-6 receptor signaling, modulated this crosstalk, attenuating macrophage activation markers and tumor cell migratory responses. Together, these findings support a model in which tumor-intrinsic PI3K signaling initiates a cytokine-dependent feed-forward circuit that reinforces macrophage reprogramming and promotes an immunosuppressive, pro-migratory microenvironment.

These mechanistic observations identify tumor-intrinsic PI3K activity as an important regulator of macrophage reprogramming in GBM. While acute PI3K inhibition in tumor cells did not produce marked changes in macrophage AKT phosphorylation, it significantly altered macrophage transcriptional programs. Notably, cytokine-related pathways, including IL-10, IL-4/IL-13, and IL-6 signaling, were enriched in macrophages exposed to PI3K-competent tumor cells and were attenuated when tumor PI3K activity was inhibited. These observations suggest that tumor-derived soluble factors, rather than direct activation of macrophage PI3K signaling, mediate this reprogramming. Importantly, the magnitude of these effects varied across GBM models, consistent with the known heterogeneity of the disease.

IL-6 emerged as a central mediator of the PI3K-dependent tumor-macrophage crosstalk identified in our model. The IL-6/STAT3 axis is known to be associated with macrophage polarization toward immunoregulatory states, upregulate PD-L1 and IDO1, promote VEGF-mediated angiogenesis, and contribute to TMZ resistance through MGMT/STAT3 signaling [38]. In GBM models, IL-6 inhibition modestly enhances T-cell infiltration and survival, indicating that targeting IL-6 can reprogram the tumor immune microenvironment [39]. In our model, IL-6 was strongly induced in both tumor cells and macrophages during coculture, and its associated signaling pathways were attenuated following PI3K inhibition. Functional blockade of IL-6 receptor signaling with tocilizumab reduced macrophage activation markers and downstream signaling responses, supporting a relevant role for IL-6 within this paracrine network. However, given the complexity of tumor-conditioned media, these effects likely reflect the combined action of multiple cytokines and signaling pathways rather than a single dominant mediator. Tocilizumab, a clinically approved IL-6R antibody, has well-established safety in humans and is already being investigated in oncology settings. Although large antibodies generally have limited penetration across an intact blood-brain barrier (BBB), and BBB disruption in GBM is often heterogeneous and overestimated by contrast enhancement, emerging small-molecule IL-6/IL-6R inhibitors with improved CNS penetration may represent an alternative strategy to target this pathway in the brain. Moreover, peripheral IL-6R blockade can modulate systemic inflammation and IL-6 trans-signaling, potentially reducing immunosuppressive programming by circulating monocytes prior to tumor infiltration [40], decreasing IL-6-mediated immunosuppression in infiltrating immune cells, and enhancing TMZ efficacy. Early translational studies have begun to explore tocilizumab combinations with radiotherapy and immunotherapy in GBM and other solid tumors, supporting feasibility [41].

Because PI3K activity lies upstream of IL-6 induction in our model, simultaneous inhibition of PI3K and IL-6 signaling may provide complementary therapeutic benefits. PI3K inhibition would suppress tumor cell-intrinsic survival pathways and inflammatory secretome programs, whereas IL-6 blockade would limit paracrine STAT3 activation in macrophages and other stromal cells. Such dual targeting provides a strong rationale for evaluating whether dual targeting produces more profound remodeling of the tumor microenvironment than either strategy alone while simultaneously enhancing sensitivity to temozolomide and immunotherapy. Consistent with this rationale, previous studies have shown that PI3K inhibition sensitizes glioma models to temozolomide [42, 43], whereas brain-penetrant PI3K inhibitors such as paxalisib and buparlisib have demonstrated clinical feasibility despite toxicity concerns [44, 45]. Nonetheless, IL-6 inhibition alone induces only modest immune activation in preclinical glioblastoma, indicating that biomarker-guided combination strategies will probably be required [39].

Several translational considerations must inform future development. First, blood-brain barrier penetration and pharmacodynamics vary across PI3K inhibitors: although buparlisib and similar agents show adequate CNS penetration and have been trialed clinically, toxicity - hyperglycemia, hepatotoxicity, and neuropsychiatric effects - remains a concern [44]. While tocilizumab is generally well-tolerated and has been safely administered in settings such as CAR T-cell therapy without compromising therapeutic efficacy, careful consideration of timing relative to cytotoxic or immunomodulatory agents remains warranted [46]. Biomarker-driven patient selection will likely be essential. Candidate markers include PI3K pathway alterations (PIK3CA/PIK3R1 mutations, PTEN loss), baseline IL-6 expression, macrophage transcriptional states (e.g., PAGE PI3K-AKT Z-scores), CIBERSORTx-derived myeloid signatures, and imaging biomarkers such as rCBV/ONCOhabitats vascular phenotypes [33].

Finally, while our data support a critical role for the PI3K-IL-6 axis in macrophage reprogramming, translation to the clinic must consider several limitations. In addition, the use of THP-1-derived macrophages may not fully recapitulate the diversity and functional states of primary human macrophages or brain-resident microglia. Further studies using recombinant cytokines or genetic approaches will be required to delineate the relative contribution of IL-6 versus other components of the tumor secretome. Pathway annotations such as “PI3K-AKT reactome” include genes regulated by multiple signaling cascades (e.g., JAK/STAT, MAPK), meaning transcriptomic changes may reflect integrated paracrine signaling rather than direct macrophage PI3K activation. More granular approaches, such as time-resolved phospho-profiling or single-cell phospho-flow, could strengthen mechanistic inference. Additionally, combinatorial regimens increase the risk of adverse events and will require meticulous dose-finding and immune monitoring [47].

In summary, our study identifies PI3K as a major regulator of inflammatory tumor-macrophage crosstalk in glioblastoma and provides evidence that this signaling axis contributes to the establishment of MES-like malignant-cell states.

## Supporting information

Uncropped westerns

## ABBREVIATIONS

AC: Astrocyte-like malignant transcriptional state
ANOVA: Analysis of variance
CCL: C-C motif chemokine ligand
CD: Cluster of differentiation
CIBERSORTx: Cell-type Identification By Estimating Relative Subsets Of RNA Transcripts
CM: Conditioned medium
CSF1: Colony-stimulating factor 1
CXCL: C-X-C motif chemokine ligand
EMT: Epithelial-to-mesenchymal transition
GBM: Glioblastoma
GO: Gene Ontology
GSEA: Gene Set Enrichment Analysis
HGF: Hepatocyte growth factor
HR: Hazard ratio
ICAM1: Intercellular adhesion molecule 1
IL: Interleukin
IPA: Ingenuity Pathway Analysis
MES: Mesenchymal transcriptional state
MRI: Magnetic resonance imaging
NF-κB: Nuclear factor kappa B
NPC: Neural progenitor-like malignant transcriptional state
OPC: Oligodendrocyte progenitor-like malignant transcriptional state
PAGE: Parametric Analysis of Gene Set Enrichment
PBS: Phosphate-buffered saline
PCR: Polymerase chain reaction
PI3K: Phosphoinositide 3-kinase
qPCR: Quantitative polymerase chain reaction
RNA-seq: RNA sequencing
scRNA-seq: Single-cell RNA sequencing
SEM: Standard error of the mean
ssGSEA: Single-sample Gene Set Enrichment Analysis
TAM: Tumor-associated macrophage
TCGA: The Cancer Genome Atlas
TGFB1: Transforming growth factor beta 1
TNF: Tumor necrosis factor
UMAP: Uniform Manifold Approximation and Projection
VCAM1: Vascular cell adhesion molecule 1
VEGFA: Vascular endothelial growth factor A

## DECLARATIONS

### Ethical Approval and Consent to participate

The study was approved by the Institutional Ethics Committee of Hospital Universitari i Politècnic La Fe, Valencia (Spain), under protocol number 2022/435.

### Consent for publication

All authors consent to the publication of this manuscript.

### Data Availability

The transcriptomic data used in this study were deposited in R2: Genomics Analysis and Visualization Platform and can be accessed through the cohort’s name Exp GBM (Macrophage tr) - FontdeMora - 16 - cpm and Exp GBM (Wortmannin tr) - FontdeMora - 24 - cpm at http://r2.amc.nl. Targeted sequencing data used to validate GBM cell lines are provided in Supplementary File 1. MRI data from TCGA IDH-wild-type GBM patients were used for ONCOhabitats analysis to define high-and low-vascular groups, which were subsequently used for evaluation of PI3K alterations and immune profiling using CIBERSORTx. Parametric Analysis of Gene Set Enrichment (PAGE) was performed within the R2 platform on macrophage-associated pathways.

### Author contributions

R.B.-P., M.H.M., and J.F.dM. conceived and designed the experiments. RNA-seq datasets were processed and analyzed by E.C. and A.E.-C., and subsequently interpreted by R.B.-P., Z.J.T., M.H.M., and J.F.dM. Z.J.T. performed the single-cell RNA-sequencing analyses. Cell culture and macrophage polarization experiments were performed by R.B.-P., M.R.-S., and N.M.-A. ddPCR experiments were performed by R.B.-P. with assistance from M.R.-S. and N.M.-A. Z.J.T. conducted PAGE and additional transcriptomic analyses using R and R2 Genomics Analysis and Visualization Platform. ONCOhabitats analyses were performed by R.B.-P., V.M.M. and J.M.G.-G. Data analysis, interpretation, and figure preparation were conducted by R.B.-P., Z.J.T., M.R.-S., and J.F.dM., with input from M.H.M. The initial manuscript was drafted by R.B.-P. and J.F.dM. All authors discussed the results and contributed to the final version of the manuscript.

## SUPPLEMENTARY FIGURES AND THEIR LEGENDS

**Supplementary Figure 1. Pharmacological confirmation of THP-1 differentiation and polarization.** THP-1 monocytes were differentiated into macrophages using PMA and subsequently polarized toward M1 (LPS + IFN-γ) or M2 (IL-4 + IL-13) phenotypes. Expression of canonical polarization markers was quantified by qPCR and/or immunoblotting. Mean ± SEM from three independent replicates per condition are shown (*p* ≤ 0.05; \**p* < 0.01; \*\**p* < 0.001; \*\*\**p* < 0.0001; one-way ANOVA). These data confirm effective pharmacological differentiation of THP-1-derived macrophages prior to their use in coculture assays with GBM cells.

**Supplementary Figure 2. Pathways differentially enriched in high-versus moderate-vascular GBM according to PAGE analysis.** Parametric Analysis of Gene Set Enrichment (PAGE) performed in R2 software comparing High Vascularity (HV) and Moderate Vascularity (MV) GBMs. Pathways with |fold change| > 1.5 and |Z-score| > 1.96 were considered significantly enriched. The tables summarize the main upregulated (left) and downregulated (right) pathways in HV tumors. The PAGE score represents the degree of enrichment, with values >1.96 corresponding approximately to *p* < 0.05, as previously described [22]. *p*-values are not shown because they are not directly comparable to standard statistical tests. These data extend the analysis shown in Figure 1D by detailing the specific PI3K-, EGFR-, PTEN-, and cytokine-related pathways contributing to the transcriptional profile of highly vascular GBMs.

**Supplementary Figure 3. PI3K inhibition with wortmannin prevents TAMs-induced GBM cellular migration.** Representative wound-healing images used for migration quantification in Supplementary Figure 5. Conditioned media were collected after 72 h from M0 macrophages cultured in transwell contact-free coculture with U87, LN-18, or LN-229 GBM cells that were either untreated or pretreated with wortmannin (200 nM, 1 h) followed by washing prior to coculture. CM from GBM cells cultured alone ± wortmannin pretreatment (Control; Wortmannin) was included as controls. Naïve GBM cells were exposed to the indicated CM conditions and imaged over time using a Leica DMi8 time-lapse microscope (10x objective). For each cell line, images at baseline and after migration are shown. Quantification of migration rates is provided in Supplementary Figure 5.

**Supplementary Figure 4. Transient PI3K inhibition by wortmannin exerts a sustained effect on AKT signaling in GBM cells.**

(A) Western blot analysis of phosphorylated (p-AKT) and total AKT protein levels. GBM cell lines were seeded in complete medium for 24 h, serum-starved for 16 h, and treated with wortmannin (200 nM) for 1 h. After drug wash-off, treated and untreated cells were stimulated with insulin (20 nM, 15 min), as indicated, to assess AKT activation as a positive control.

(B) Time-course analysis of p-AKT levels following wortmannin wash-off showing prolonged inhibition of PI3K signaling up to 72 h.

(C) Western blot analysis of cleaved and total caspase-3 during the wortmannin time-course experiment to evaluate apoptotic activation. Data are representative of three independent experiments.

**Supplementary Figure 5. An NF-κB inflammatory program and the four-state classification recapitulate the mesenchymal-communication association.**

(A) UMAP representing all 7,930 cells colored by cell-type (malignant, macrophage, T cell, oligodendrocyte); non-malignant lineages were identified by the Neftel marker-set (mean marker expression > 4) and excluded from downstream analyses.

(B) UMAP of the malignant cells colored by the NF-κB inflammatory program score.

(C) Within-tumor Spearman correlations (ρ) between the MES-like score and the NF-κB score, one point per patient; point size denotes malignant-cell number. The subtitle reports the median within-tumor ρ and the linear mixed-model fixed-effect p-value.

(D) Tumor-level association between the percentage of MES-like malignant cells and the mean NF-κB score; one point per tumor, point size denotes malignant-cell number, line and shaded band show the linear fit with 95% confidence interval, and the subtitle reports the Spearman ρ and p-value.

(E-G) Tumor-level associations between the percentage of cells in each remaining Neftel state and the mean NF-κB score (left) and mean tumor-macrophage communication signature score (right), for the astrocyte-like (AC; E), oligodendrocyte-progenitor-like (OPC; F) and neural-progenitor-like (NPC; G) states. As the four state fractions are compositionally constrained to sum to 100%, these non-MES associations reflect relative rather than independent effects. Each subtitle reports the Spearman ρ and p-value.

**Supplementary Figure 6. Conditioned media from GBM-macrophage cocultures enhances tumor cell migration through PI3K-dependent mechanisms.** GBM cells were exposed for 12 h to CM collected from GBM-macrophage cocultures (3 days). Prior to coculture, GBM cells were pre-treated with wortmannin (200 nM, 1 h) and washed twice to remove the inhibitor. Quantification of migration rates for U87 (A), LN18 (B), and LN229 (C) cells. CM from cocultures markedly increased migration in all lines, whereas CM generated from wortmannin-pre-treated GBM cells failed to enhance migration. Data represent mean ± SEM of triplicates from three independent experiments; statistical significance was determined by one-way ANOVA with Tukey multiple comparisons test (*p* ≤ 0.05; \**p* < 0.01; \*\**p* < 0.001; \*\*\**p* < 0.0001). Where indicated, direct wortmannin treatment of GBM cells (without CM) reduced migration, confirming that PI3K signaling intrinsically contributes to motility.

**Supplementary Figure 7. PAGE analysis of macrophage transcriptional responses following coculture with GBM cells.**

(A) PAGE (Parametric Analysis of Gene-set Enrichment) was performed in the R2 Genomics platform using RNA-seq data from macrophages cocultured with GBM cell lines (U87-MG, LN-18, LN-229) compared with unpolarized M0 macrophages. Cocultured macrophages exhibited strong enrichment of IL-4/IL-13 signaling, IL-10 signaling, interferon-γ-related pathways, and IL-1-associated pathways (indicated by *), consistent with the induction of a mixed but predominantly M2-like macrophage phenotype.

(B) PAGE analysis comparing macrophages exposed to GBM cells pre-treated with wortmannin versus M0 controls revealed no enrichment in these cytokine-driven Reactome pathways. The absence of these signatures supports that PI3K activity in GBM cells is required for the induction of M2-associated transcriptional programs in macrophages.

(C) Summary table of PAGE rankings and PAGE scores for the five key cytokine pathways (IL-10, IL-4/IL-13, IFN-γ, IL-1, IL-6). All five pathways showed marked decreases in both PAGE ranking and PAGE score when glioblastoma cells were pre-treated with wortmannin. These quantitative reductions demonstrate that PI3K inhibition in glioblastoma cells substantially blunts the induction of macrophage cytokine-driven programs, further supporting the role of tumor PI3K signaling in promoting an M2-like tumor-associated macrophage phenotype.

**Supplementary File 1. Targeted DNA sequencing of GBM cell lines.** Summary of somatic variants detected in LN-18, LN-229, and U-87 using the Oncomine Comprehensive Assay Plus. The table includes pathogenic or likely pathogenic mutations, allele frequencies, and affected pathways. Notably, LN-18 harbors the PIK3CB p.E1051K mutation (allele frequency 100%), U87-MG carries a PTEN splice-site variant (NM_000314.8) at 100% allele frequency, and LN-229 shows no pathogenic mutations in PI3K pathway genes, confirming wild-type PI3K/PTEN status.

**Supplementary Table 1. Characteristics of malignant cells retained for single-cell analyses.**

**Supplementary Table 2. Patient-level summary of malignant-cell state composition and inflammatory program scores.**

**Supplementary Table 3. Macrophages Subset Gene Signatures.**

