## Supplementary figures and images for "PI3K signaling promotes inflammatory tumor-macrophage crosstalk associated with mesenchymal glioblastoma"

### Supplementary Figure 1

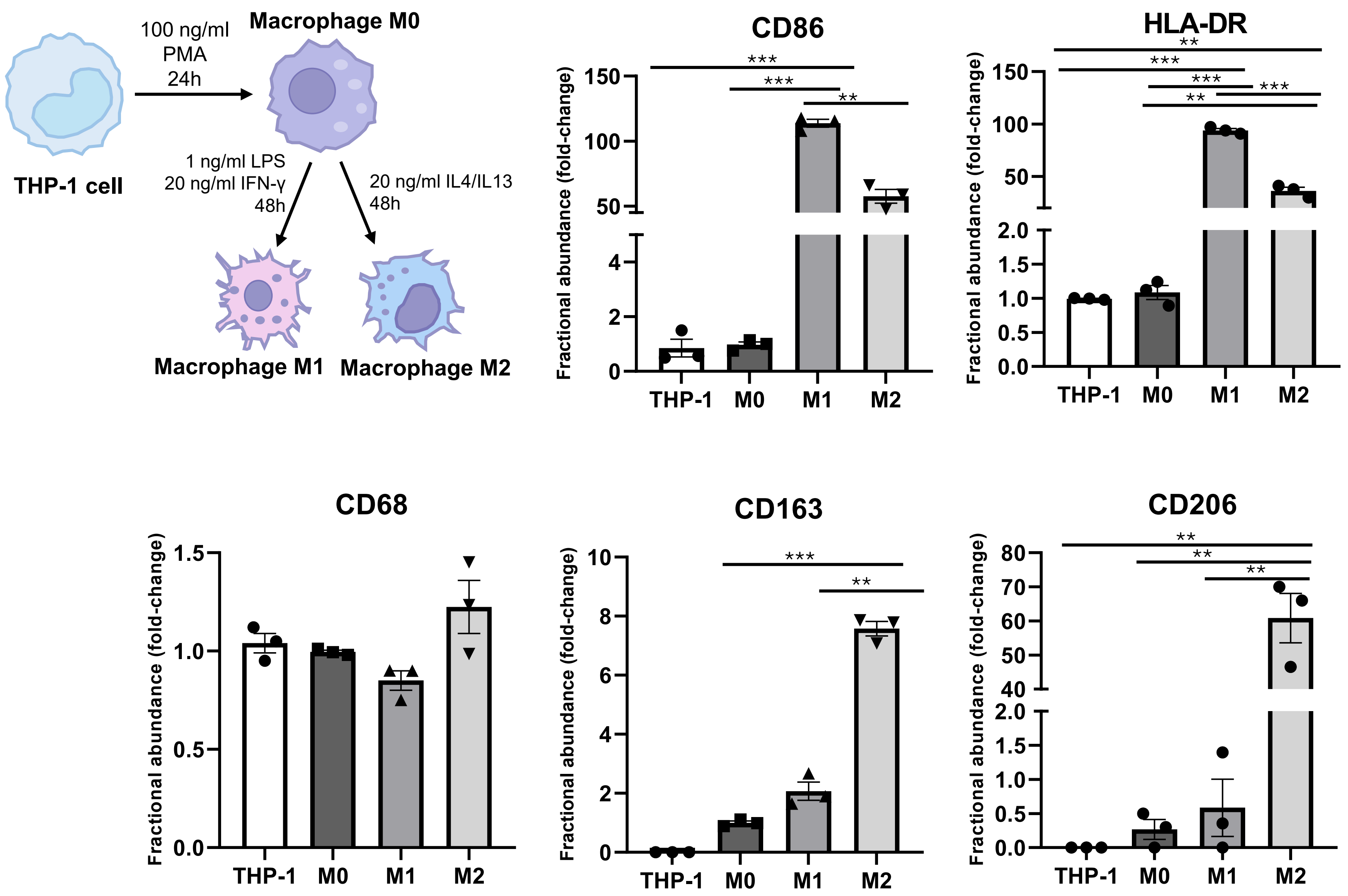

### Supplementary Figure 3

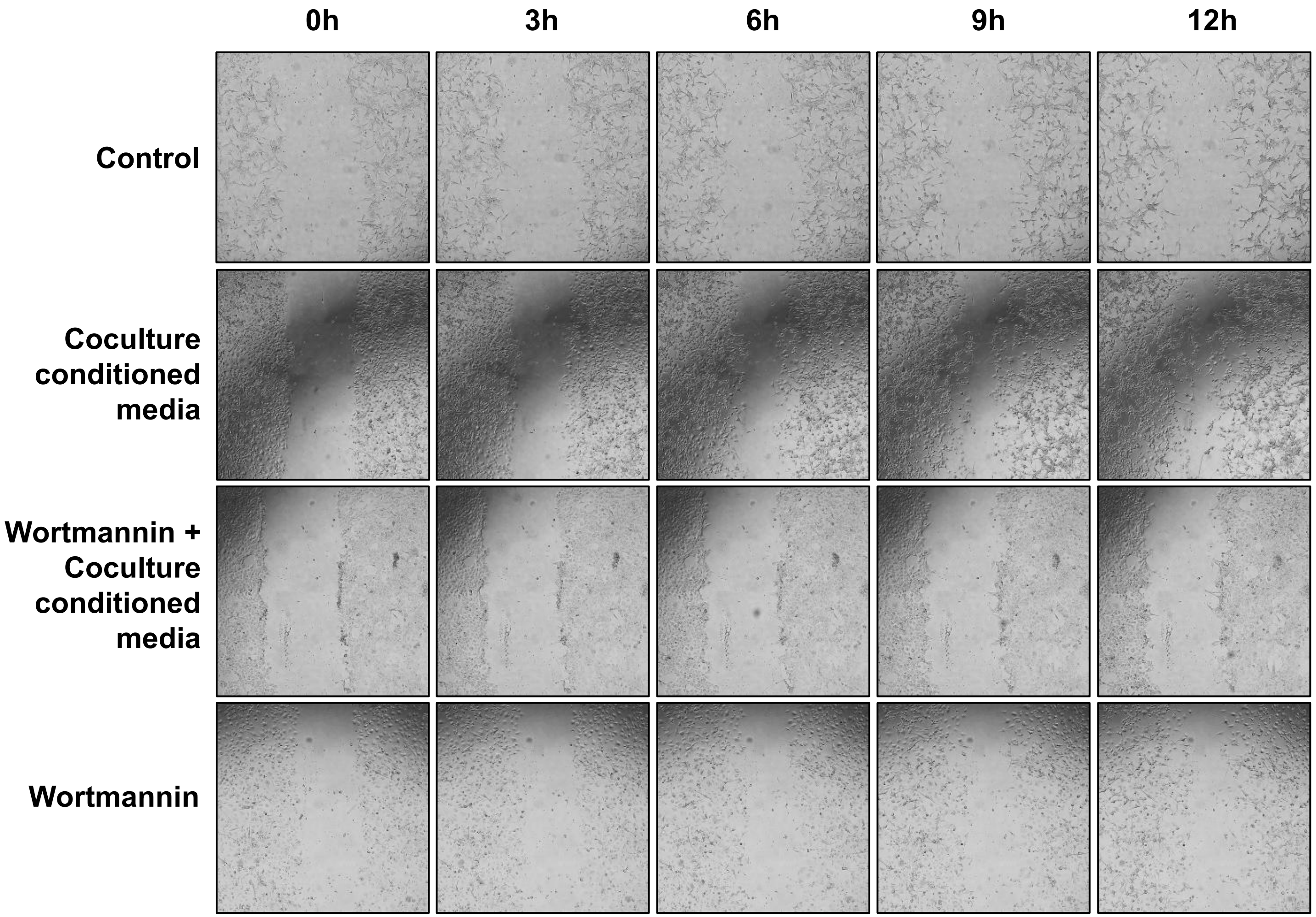

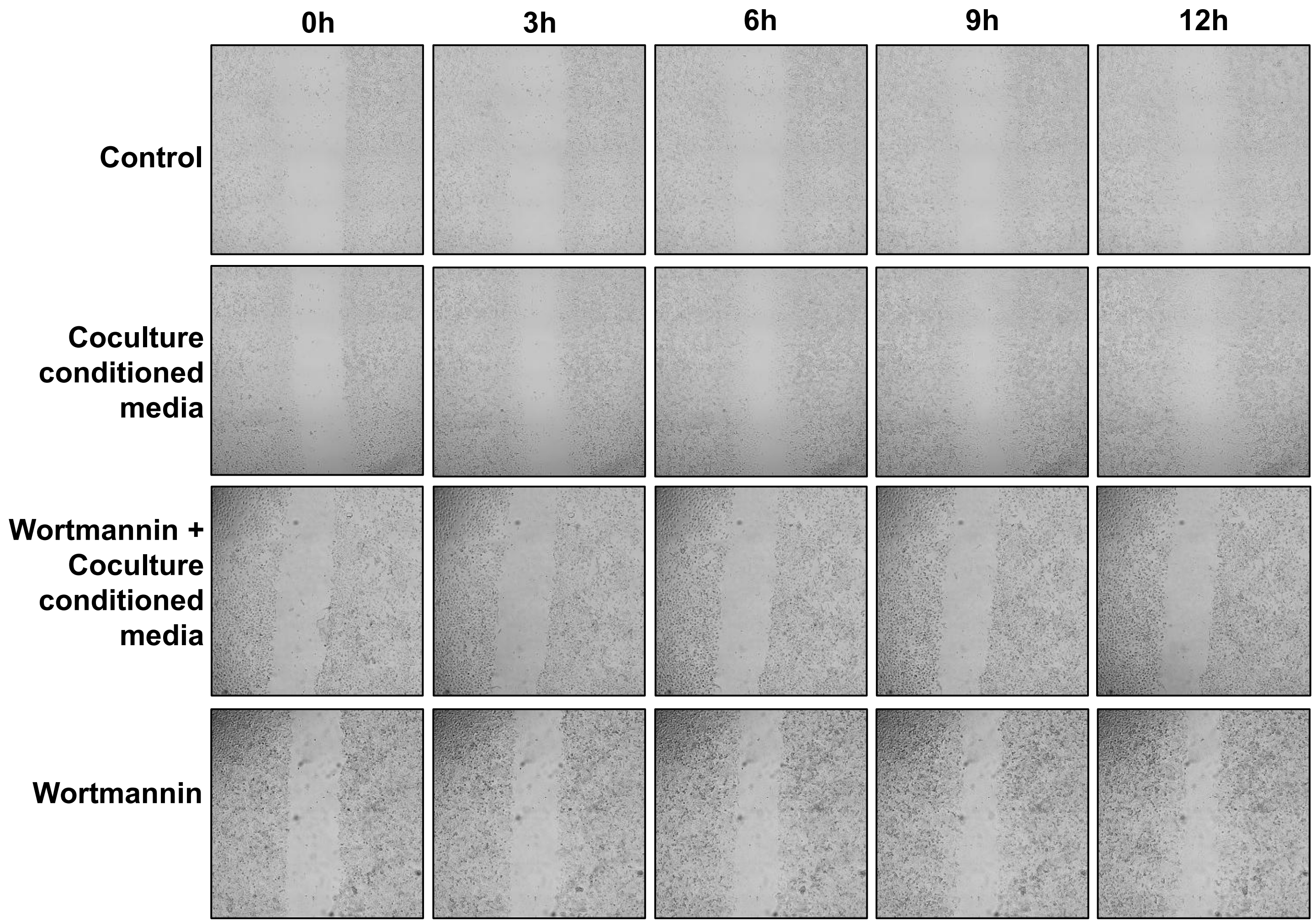

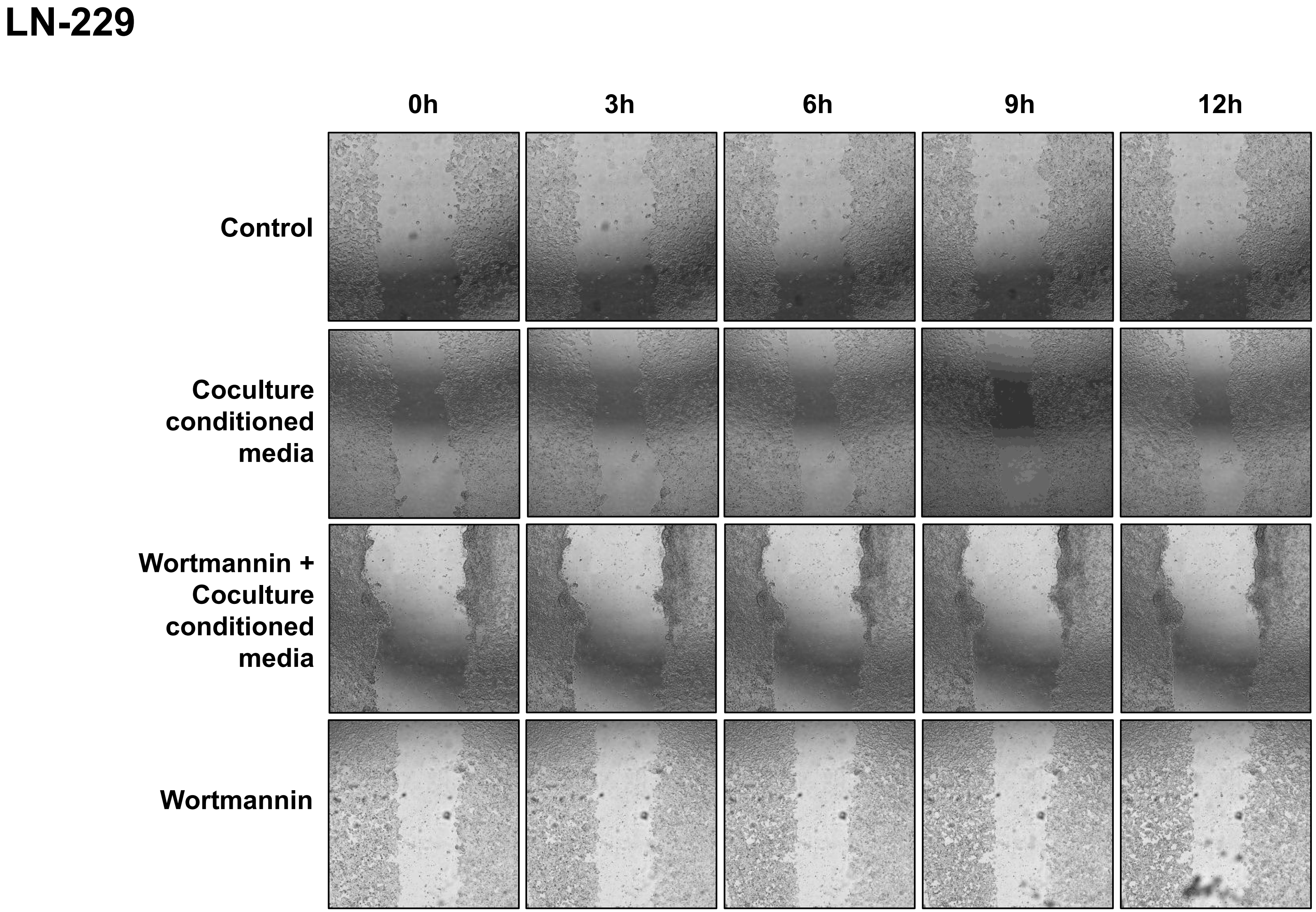

### Supplementary Figure 4

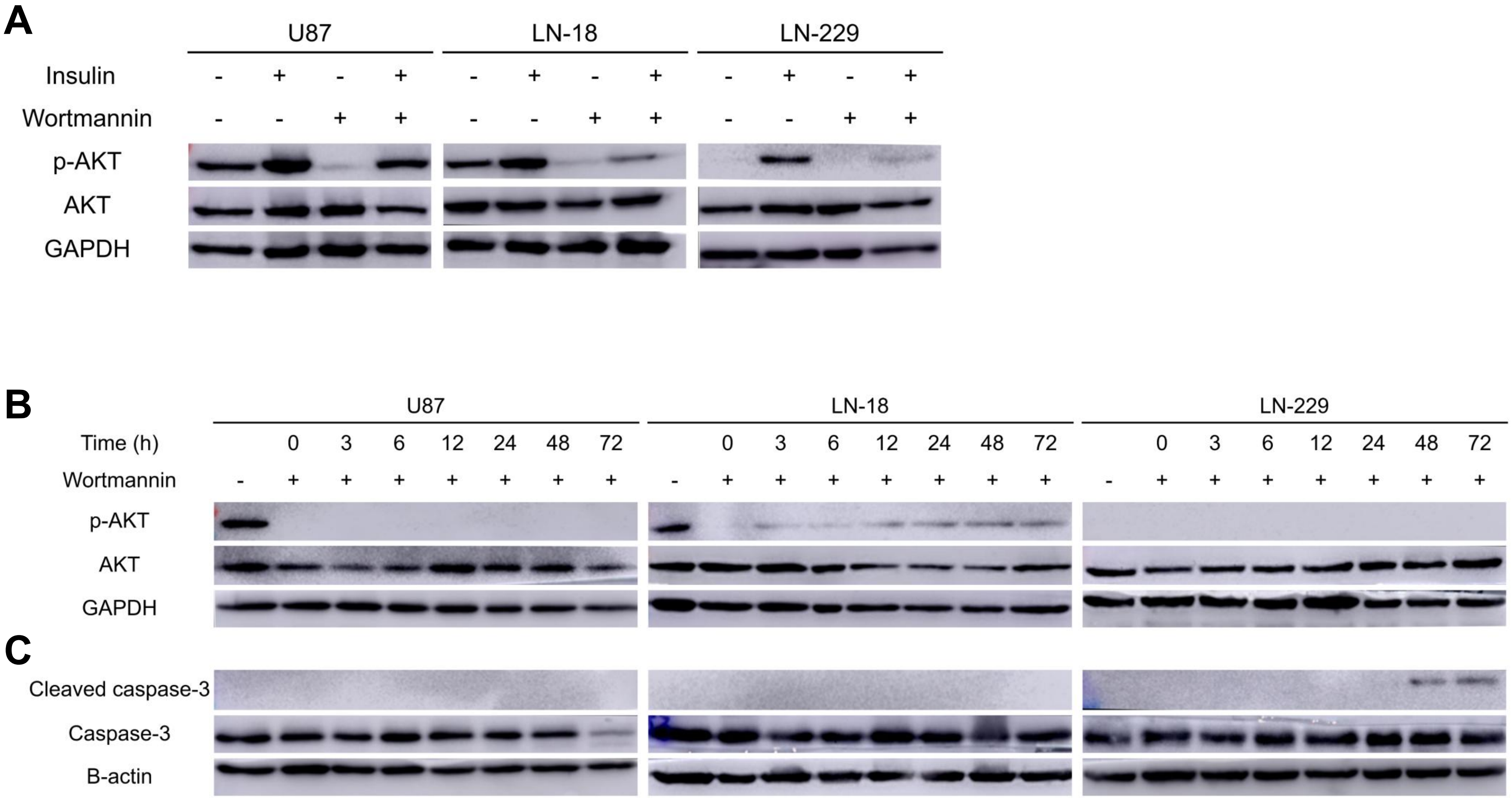

### Supplementary Figure 5

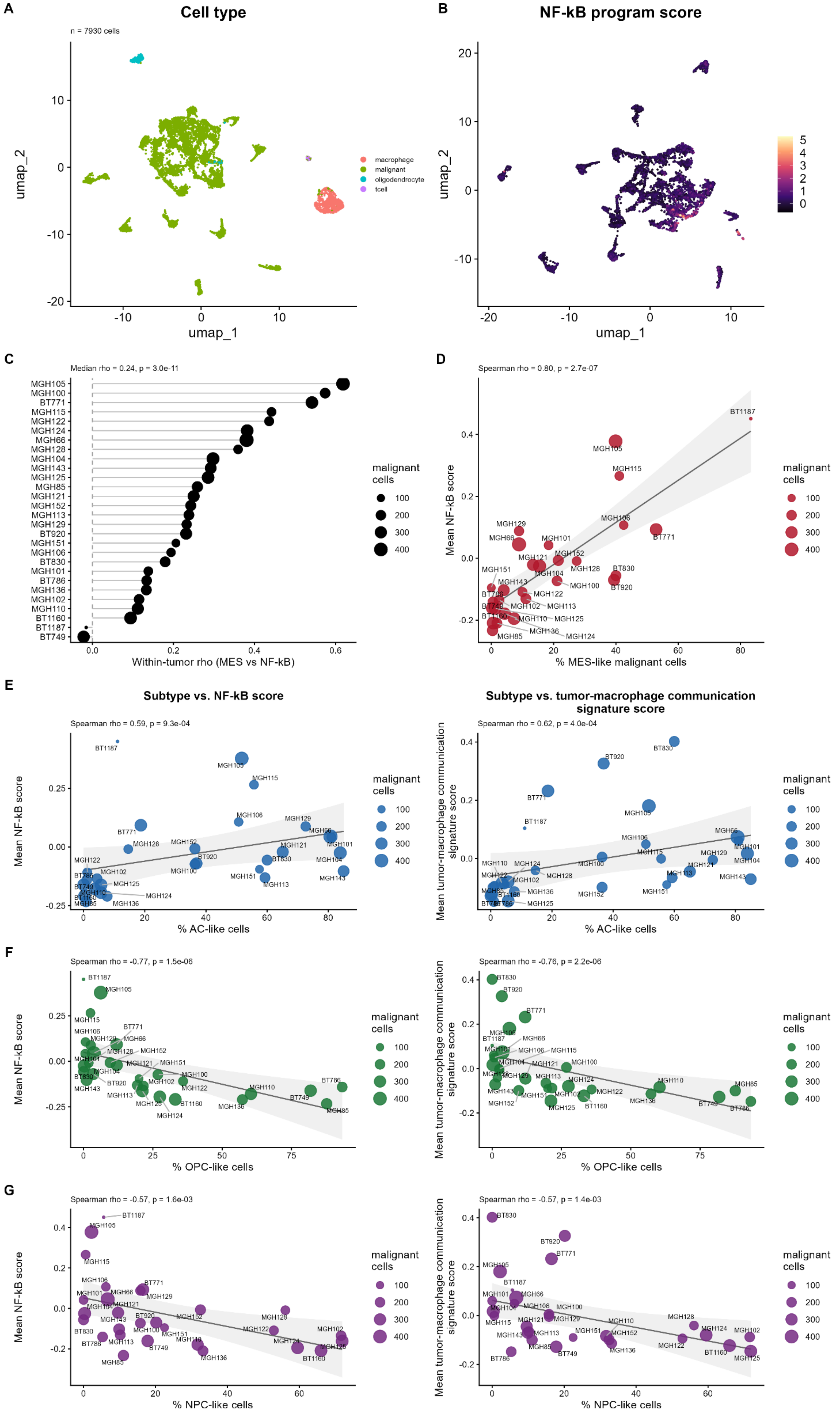

### Supplementary Figure 6

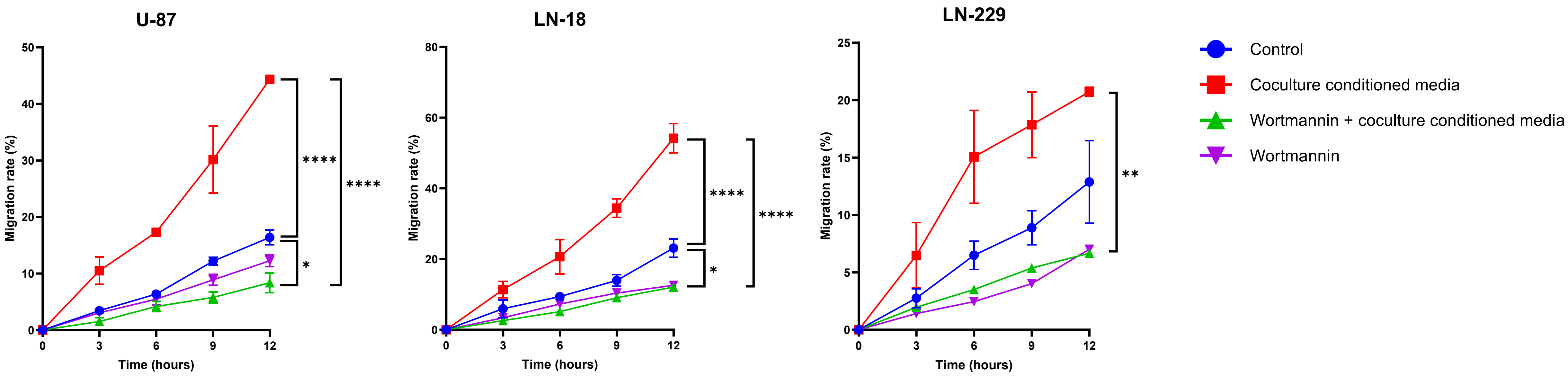

### Uncropped westerns

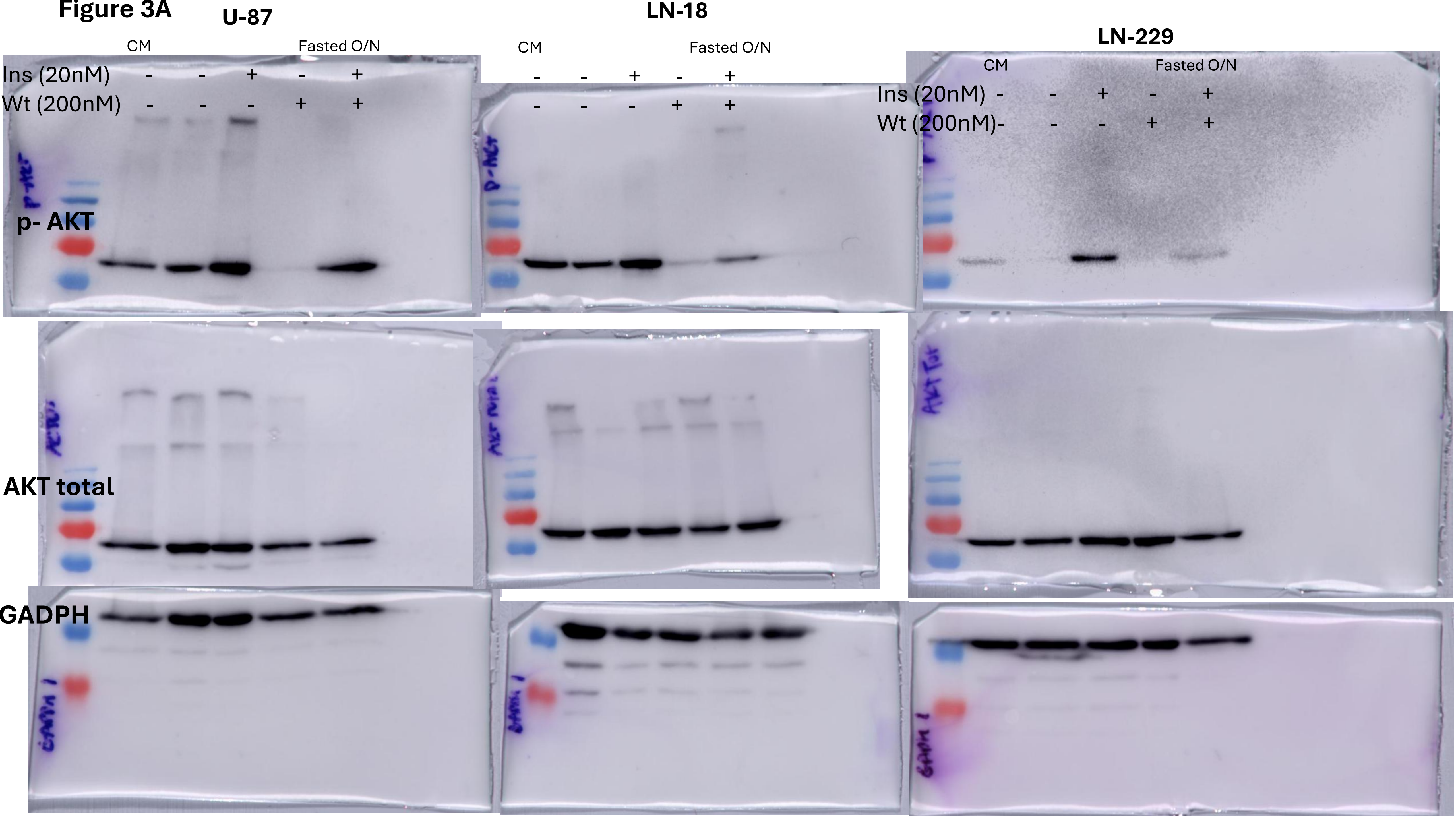

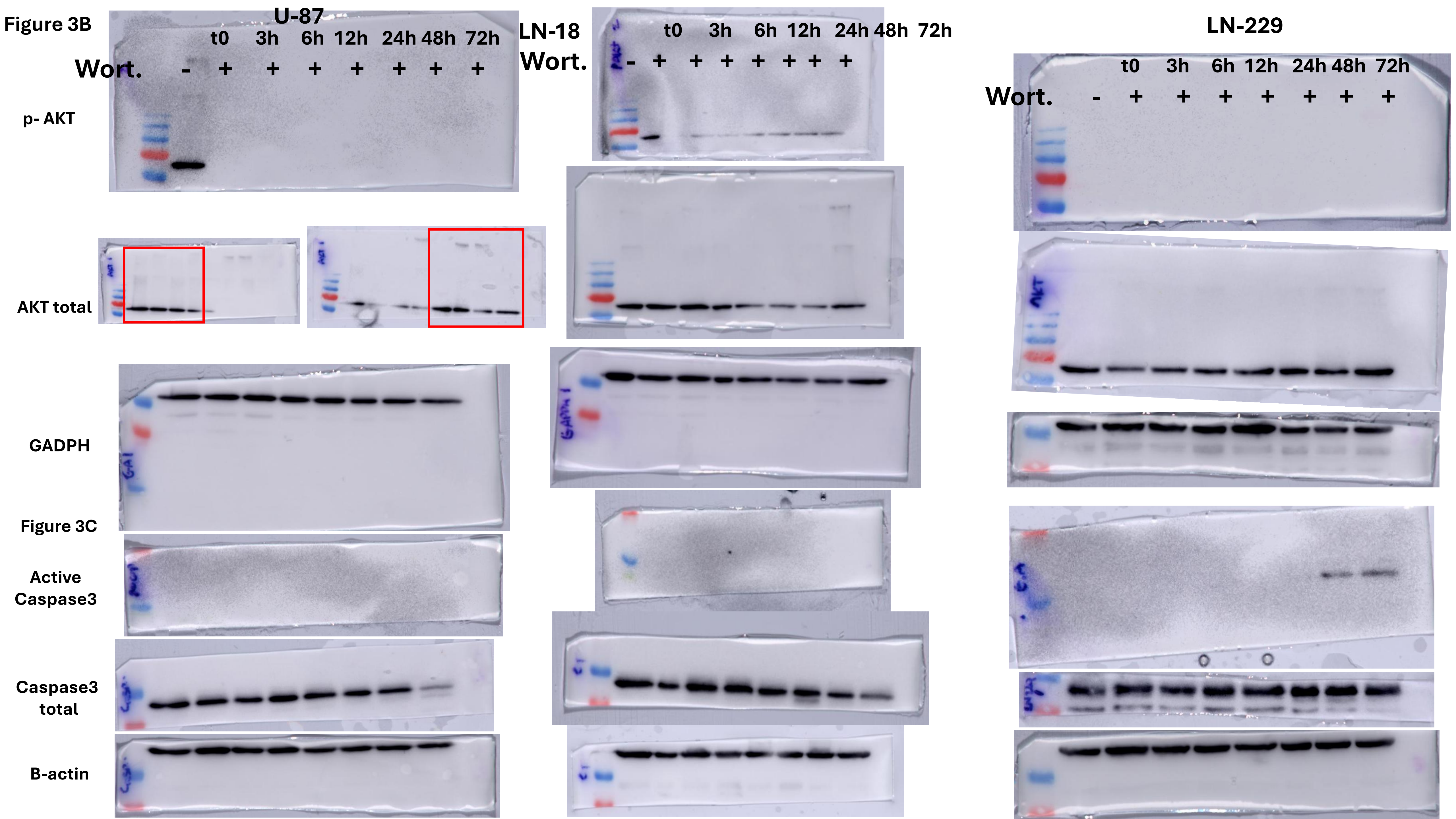

Figure 7A

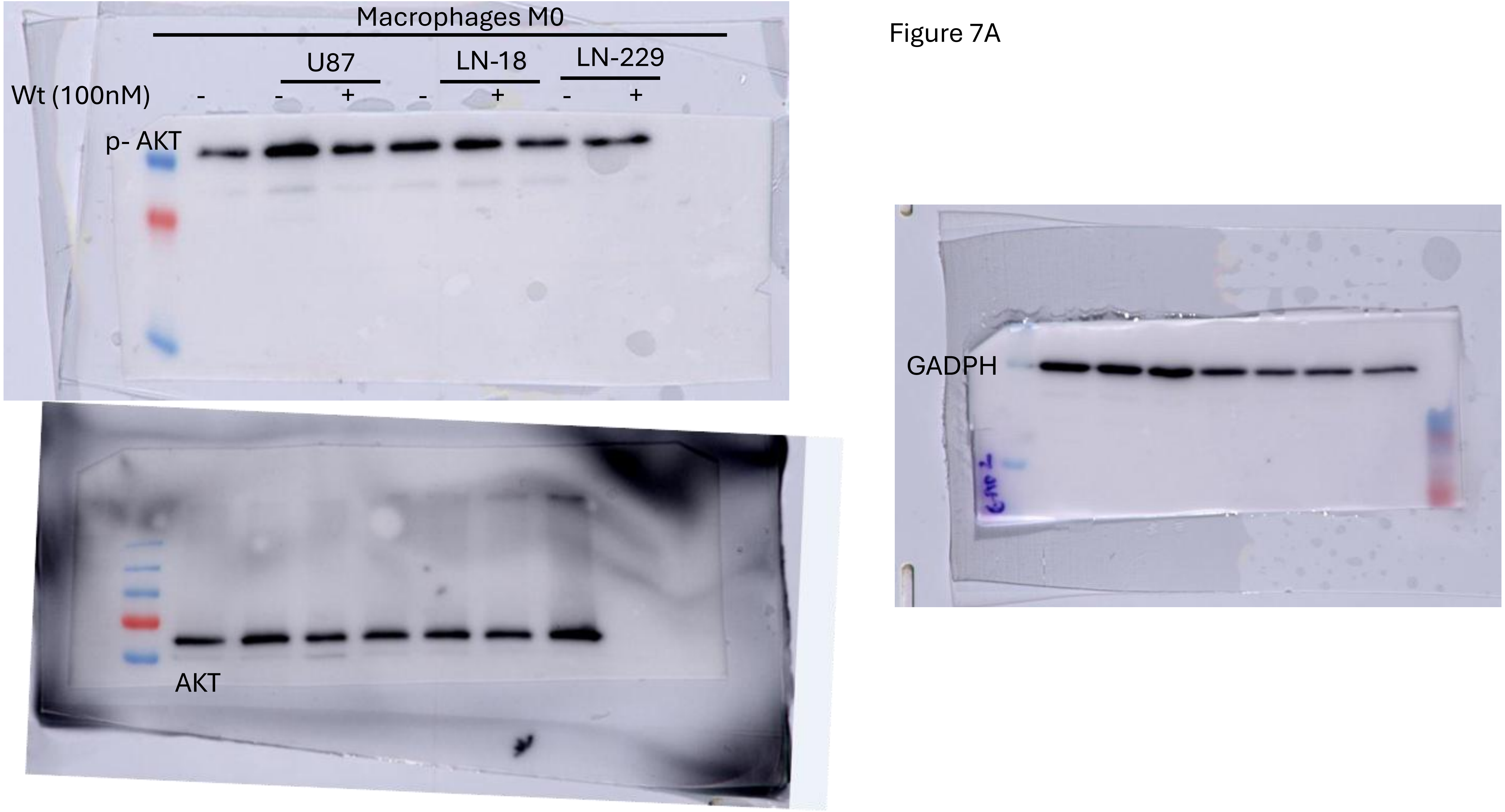

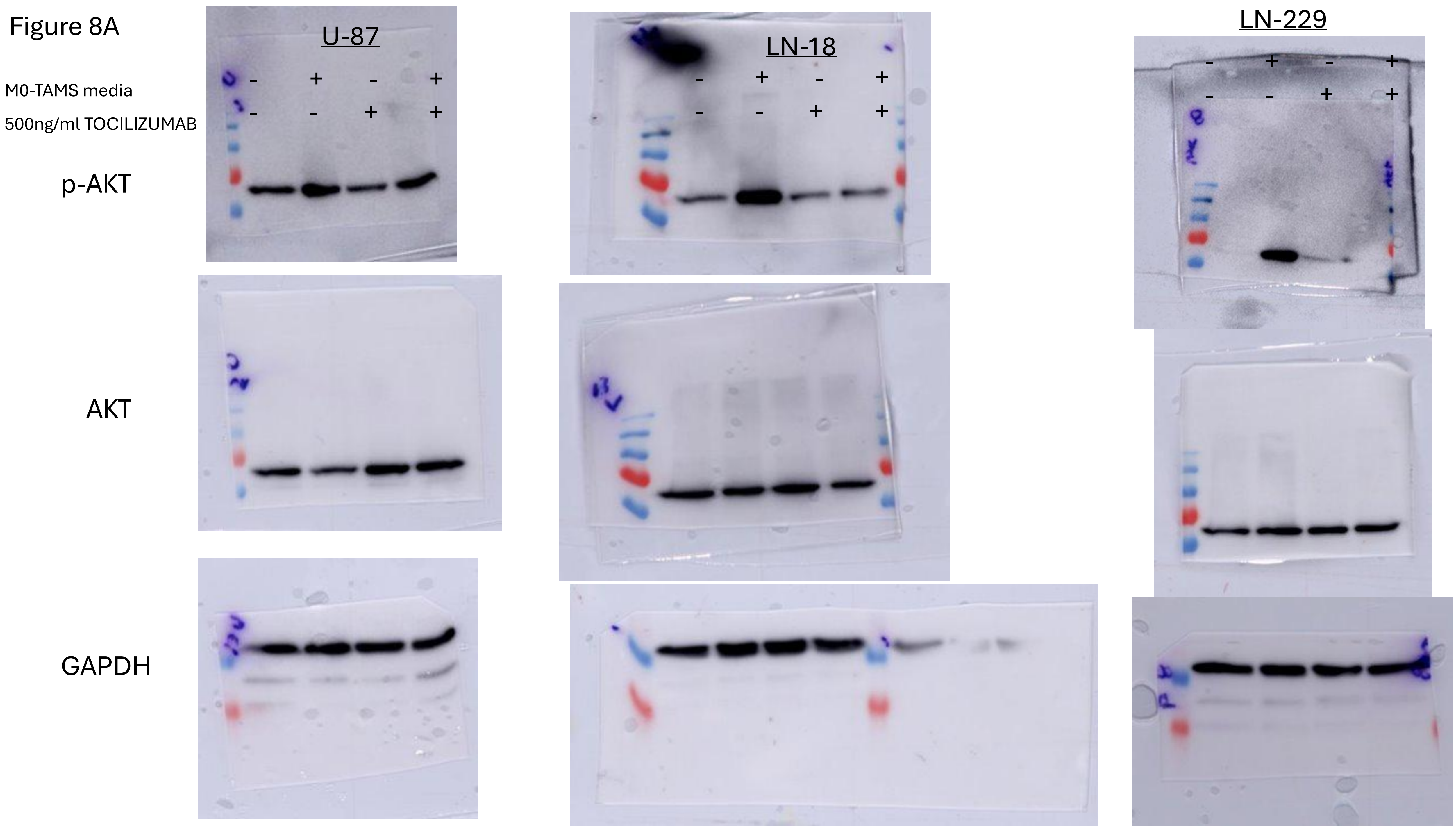
