## Supplementary Figure 2 for "PI3K signaling promotes inflammatory tumor-macrophage crosstalk associated with mesenchymal glioblastoma"

### **Upregulated pathways** in High vascular group

| Gene Set | PAGE score |
|---|---|
| **VERHAAK_GLIOBLASTOMA_MESENCHYMAL (214)** | 5.492 |
| **REACTOME_INNATE_IMMUNE_SYSTEM (864)** | 4.614 |
| **REACTOME_INTERLEUKIN_10_SIGNALING (42)** | 4.395 |
| **REACTOME_INTERLEUKIN_4_AND_INTERLEUKIN_13_SIGNALING (103)** | 3.461 |
| **REACTOME_CONSTITUTIVE_SIGNALING_BY_ABERRANT_PI3K_IN_CANCER (67)** | 3.176 |
| **REACTOME_CYTOKINE_SIGNALING_IN_IMMUNE_SYSTEM (763)** | 3.170 |
| **REACTOME_PI_3K_CASCADE_FGFR2 (22)** | 2.629 |
| **REACTOME_PI_3K_CASCADE_FGFR4 (18)** | 2.602 |

### **Downregulated pathways** in High vascular group

| Gene Set | PAGE score |
|---|---|
| **KOBAYASHI_EGFR_SIGNALING_24HR_DN (244)** | -13.816 |
| **CROONQUIST_IL6_DEPRIVATION_DN (95)** | -7.870 |
| **REACTOME_PTEN_REGULATION (119)** | -5.473 |
| **REACTOME_REGULATION_OF_PTEN_STABILITY_AND_ACTIVITY (65)** | -4.377 |
| **REACTOME_REGULATION_OF_PTEN_GENE_TRANSCRIPTION (52)** | -3.043 |
