## Supplementary Figure 7 for "PI3K signaling promotes inflammatory tumor-macrophage crosstalk associated with mesenchymal glioblastoma"

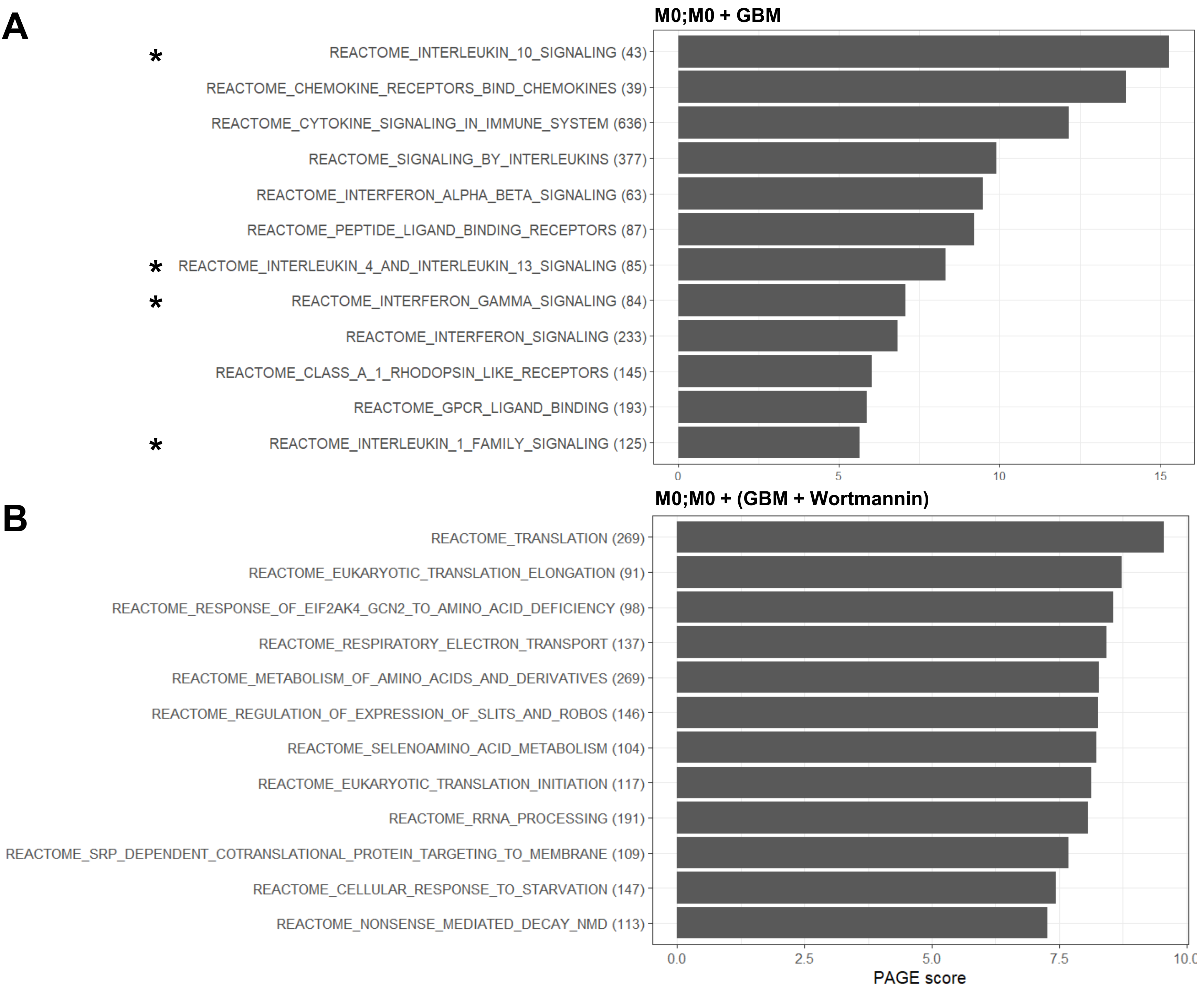


**C**

| | M0;M0+GBM | | M0;M0+(GBM+ Wortmannin) | |
|---|---|---|---|---|
| Reactome pathway | PAGE Rank | PAGE Score | PAGE Rank | PAGE Score |
| **IL-10** | 1 | 15.28 | 16 | 6.434 |
| **IL-4/IL-13** | 7 | 8.327 | 85 | 2.998 |
| **IFNγ** | 8 | 7.076 | 124 | 2.386 n.s. |
| **IL-1** | 12 | 5.641 | 98 | 2.798 |
| **IL-6** | 47 | 3.058 | 577 | -0.078 n.s. |

n.s. not significant
